# Microbial communities in tropical soils are highly resilient to fluctuating redox conditions

**DOI:** 10.64898/2026.07.27.741047

**Authors:** Ashley Campbell, Ikaia Leleiwi, Amrita Bhattacharyya, Jeffrey Kimbrel, Yang Lin, Malak M. Tfaily, Allison Thompson, Rosalie Chu, Gareth Trubl, Whendee L. Silver, Ljiljana Paša-Tolić, Peter Nico, Jennifer Pett-Ridge

## Abstract

Many wet tropical soils alternate frequently between fully oxygenated and anaerobic conditions, constraining the terminal electron acceptors available for microbial metabolism, and the mineral-organic matter interactions that regulate many aspects of soil carbon (C) cycling. However, it is still unclear how fluctuating soil redox conditions influence the microbial community composition and if microbially mediated C flux is sensitive to extended oxic or anoxic periods like those observed during drought and flooding respectively. Using a 44-day redox manipulation of tropical soils that experience daily-to-weekly oxygen (O_2_) fluctuations in the field, we measured how different redox regimes shape soil biogeochemistry and microbial and metabolite composition. Replicate microcosms were exposed to four treatments (static oxic, static anoxic, high frequency fluctuation (4 day oxic/4 day anoxic), or low frequency fluctuation (8 day oxic/4 day anoxic)) regimes, and harvested for microbial, metabolite, carbon dioxide (CO_2_) flux, and biogeochemical assays at multiple timepoints. Oxic and fluctuating redox conditions caused the microbial community to shift in a manner correlated with soil iron content and directly orthogonal to communities from anoxic soils. The identity of both iron oxidizers and iron reducers was distinct in static anoxic soils but was resilient to redox fluctuation and prolonged O_2_ exposure. The total amount of CO_2_ respired was similar across all four redox regimes. Water-extractable organic matter composition was distinct across redox treatments, with anoxic soils accumulating higher levels of carbohydrate-, proteins-, amino sugar-, and lignin-like compounds consistent with reduced enzymatic decomposition and release of mineral-associated organic matter via iron reduction, while oxic soils showed elevated lipid- and unsaturated hydrocarbon-like compounds indicative of greater microbial biomass turnover. The microbial community adapted to dynamic redox conditions and the results substantiate cycling of distinct C compounds under varying redox conditions resulting from varying bioavailability (driven by mineral-OM dynamics) and/or shifted microbial metabolism.

## Introduction

Soil environmental properties such as pH and soil organic C (quantity and quality) are known drivers of microbial community composition [1–3], but an emerging consensus suggests that soil [O_2_] and redox status have an equally large influence on the structure and metabolic capacity of soil microbial communities [4, 5]. As such, studies have shown the characteristic redox oscillation in soils to drive microbial community structure [1, 2, 5, 6]. The soil properties that govern microbial community structure also play an important role in regulating microbial processes [3, 4]. Greenhouse gas fluxes, soil nutrient availability (phosphorus (P), nitrogen (N), potassium (K)), and carbon (C) cycling are sensitive to climate change and are directly or indirectly regulated by soil moisture and redox. Soil moisture regulates many redox-sensitive aspects of microbial metabolism, controlling oxygen (O_2_) and solute diffusion, availability of terminal electron acceptors, and fermentation and aerobic versus anaerobic respiratory processes.

Wet tropical soils can alternate frequently between fully oxygenated and anaerobic conditions, constraining both the metabolism of tropical soil microorganisms, and the mineral-organic matter relationships that regulate many aspects of soil C cycling. We have little understanding of how climate change will affect the community composition and consequently microbial functions in tropical ecosystems. Tropical forests are predicted to experience a 2–5°C temperature increase and substantial differences in the amount and timing of rainfall in the coming half century [7, 8]. Precipitation and temperature patterns have been shown to be closely linked with soil O_2_ concentrations [9–11]. Humid tropical forests often experience a periodic O_2_ limitation in surface soils at scales of microsites to entire soil profiles [3, 9, 11–15]. While the impact of O_2_ limitation has been well studied in wetland systems [16], the impact of O_2_ limitation on soil C storage in terrestrial soils has received less attention. For example, along a precipitation gradient in Hawaii an increase in soil C stocks was observed with decreasing redox potential, suggesting that reducing conditions promoted soil C accumulation [13]. Over the past decades, the humid, tropical Luquillo Experimental Forest (LEF) has experienced uncharacteristically severe droughts during the dry season [17, 18]. A reduction in soil water availability and an increase in the intensity and frequency of drought periods can lead to reduced decomposition and microbial growth and cause changes in the microbial community structure [19, 20]. Interestingly, in tests where LEF microbial communities have been experimentally pushed outside of their normal oscillating redox regimes and forced to tolerate a consistently aerobic or anaerobic environment—they lose biomass and taxonomic richness, rates of N, P, and Fe transformations decline, and decreased degradation of some types of complex macromolecular C [21–24]. However, the community ecology mechanisms that underlie these effects are virtually unknown.

Prior work assessing drought impacts on soil microbial community composition indicate active community membership is most affected by reduced moisture conditions and even small reductions in water potential can influence community functional potential and microbial activity [20, 25]. The O_2_ concentration in tropical soil is affected by both moisture and topography. Multiple studies at LEF show soil O_2_ concentrations inversely correlate with soil moisture conditions, particularly in valley regions, and that periods of drought increase soil carbon dioxide emissions yet decrease methane emissions[11, 17]. Iron(III) oxide-hydroxide minerals influence many biogeochemical processes through the sorption of nutrients which subsequently constitute a substantial fraction of soil organic C oxidation via microbial reductive dissolution of iron(III) oxide-hydroxides [26, 27]. Association of soil organic C with minerals is considered one of the major factors governing the soil organic C residence time in soil [28, 29]. Energy-rich organic compounds, such as carbohydrates, which are readily respired by microorganisms, have been observed to persist in soils for longer than expected based on their thermodynamic properties, potentially due to decreased bioavailability from associations with minerals [30–32]. The oxidation state of iron in tropical soils influences phosphorus and C mobility, shaping not only microbial community structure but also function [11, 33]. While it is clear elemental cycling and C persistence in soil depends on environmental factors like soil oxidation state, how bacterial and fungal membership change in response to soil redox and how their interactions with critical elements vary with redox has yet to be expounded. Resolving this question is important because climate-driven shifts in rainfall patterns are expected to alter not only soil moisture but also the temporal dynamics of O_2_ availability that govern microbial metabolism and ecosystem C storage. We assessed the effects of redox periodicity on the bacterial and fungal community and its relation to C transformation (via metabolite analysis) in surface soils from the LEF in a temporally resolved redox oscillation study. Given the inherently dynamic nature of the soil redox conditions in the LEF soils, we hypothesize that static redox conditions would alter the soil microbial community relative to the native soil, while the microbial community in the fluctuating redox treatments will more closely resemble that of the native soil. We hypothesize that each redox condition will exhibit distinct types of metabolism (e.g., heterotrophic, fermentative) which will result in significantly different soil metabolic profiles. Finally, we hypothesize that the total C respiration will be lower in the fluctuating redox treatments due to higher expected energy demands necessary to accommodate changing redox conditions.

## Methods

### Field site, sampling and soil characteristics

14.5 kilos of soil were collected from a mid-slope position at the El Verde field site in the Luquillo Experimental Forest (LEF), Puerto Rico, part of the NSF-sponsored Long-term Ecological Research (LTER) program. Soil at this site are volcanoclastic Oxisols in a Tabunuco forest that receives 2000 to 6000mm annual rainfall [34–36]. LEF soil is clay-rich with 57 +- 3% sand, 23 +- 1% silt, and 20 +- 3% clay [17, 37] with a native pH of ∼5 and enriched in short-range ordered iron mineral phases [38]. Samples were collected within a 10 x 10m area (N 18.3202, W 065.81721) by first removing the litter layer from the soil surface and then collecting soil with a sterile bulk density corer and shovel to a 10cm depth. Collected soils were transported from the field at ambient temperature and moisture and shipped overnight to Lawrence Livermore National Laboratory for immediate processing. There, the soil was lightly homogenized by removing macrofauna, roots, plant matter, and rocks. A subsample was stored at −80°C to provide a baseline measurement for ‘field’ conditions and microbial communities. Another subset was air dried.

Soil C and N content were analyzed at the University of California, Berkeley using an isotope ratio mass spectrometer. Soils contained approximately 58mg total C g soil^-1^, 4mg N g soil^-1^, and a C:N of 14.5. Field condition soil water content (43%) and water holding capacity (WHC; 0.55g), gravimetric water content of field moist soil (0.76), gravimetric content at 100% WHC (1.21), and the water content of field moist soil as a percent of WHC (63% WHC) were measured.

### Experimental laboratory incubation

Homogenized soil was divided into microcosms for a well replicated, temporally resolved incubation experiment. Replicated microcosms (n = 180) were divided between four redox treatments described in ‘treatments and harvest’ below (Fig. 1). Microcosms were 485mL Mason jars with modified lids for sampling and flow-through gas exchange. Approximately 20g dry soil weight of homogenized soil was weighed directly into microcosm jars. Soils were acclimated in the dark during a pre-incubation period of 16 days under a four-day oxic, four-day anoxic fluctuating redox regime (Fig. 1, ‘pretreatment’ gray area of timeline), which in prior work was shown to best resemble natural redox oscillation conditions for these soils [11, 21]. Soils were pre-incubated to stabilize soil respiration resulting from sampling and homogenization disturbance. After the ‘pretreatment’ we harvested a triplicate set of microcosms under the anoxic (N_2_) head space, switched the headspace to oxic (air, 20% O_2_) conditions, and harvested triplicate microcosms 30 minutes and 3 hours after complete turnover of the headspace following the switch. These nine jars serve as a baseline (’Time Zero’; ‘T_0_’).

**Figure 1.**
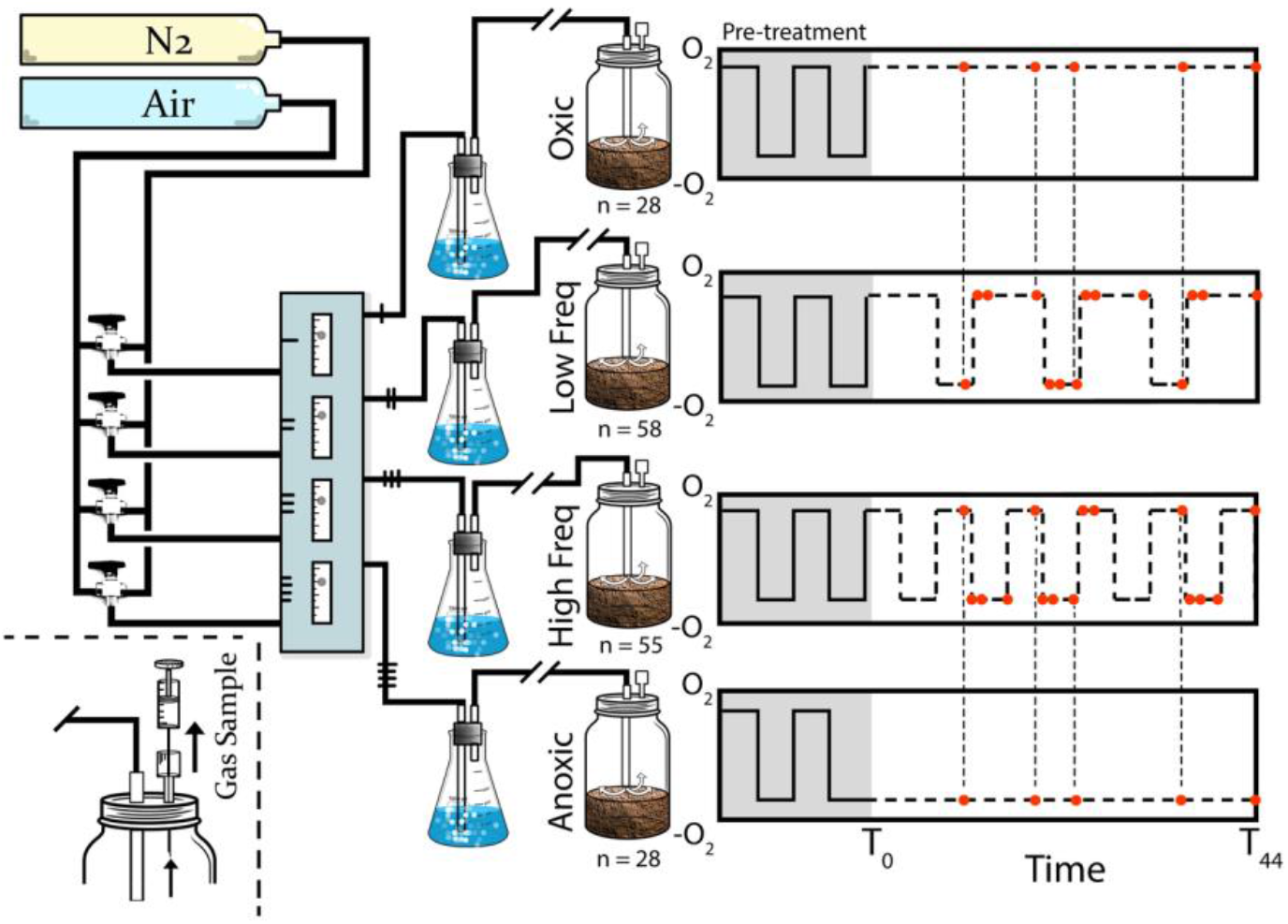
Experimental design of soil microcosm redox incubation of a wet tropical soil from the Luquillo Experimental Forest, Puerto Rico, including gas flow-through system and sampling scheme. Oxic and anoxic soil conditions were imposed by changing microcosm headspace by flushing with either N_2_ or air. After collection from the field site and homogenization, soils were weighed into jars and pre-incubated for two weeks under a 4 days oxic/ 4 days anoxic regime, then amended with plant biomass (representing 7.5% soil C) at T_0_. Four different redox treatments were applied to individual microcosms (n = 169), static oxic, static anoxic, low frequency fluctuating (8 days oxic / 4 days anoxic), and high frequency fluctuating (4 days oxic / 4 days anoxic). Microcosms were destructively harvested at multiple time points during the 44-day incubation (indicated by red dots) in replicates of at least 3. Headspace gas was repeatedly sampled (non-destructively) from the set of jars harvested at the final timepoint.

The remaining 169 microcosms were amended at T_0_ (Fig 1) with 180mg of finely ground plant biomass (PB) at a rate of 9mg g soil^-1^, (Isolife #U-61789, Wageningen, Netherlands), representing 7.5% of soil C content. Plant biomass was added to the surface of the soil using a fine sifter for even dispersal across the soil surface. Water was added to achieve 50% water content (∼3.5mL) using a glass bottle mister and aided in PB incorporation to soil. All microcosms were weighed and checked repeatedly throughout the experiment to ensure soils maintained 50% water content. No water loss was detected from microcosms throughout the experiment; air and N_2_ gases were bubbled through sterile water before reaching the microcosm headspace to prevent drying out the soils (Fig. 1). A handheld flowmeter (Proflow 6000, Restek Corp, Bellefonte, PA) was used to check gas efflux rate to ensure all microcosms experienced the same flow rate (3mL min^-1^); adjustments were made as needed using upstream flow controllers (Series RM Polycarbonate Flowmeter model RMA-150-SSV, Dwyer, Michigan City, IN). Complete replacement of the microcosm headspace for each redox switch (oxic to anoxic or vice versa) took 30 minutes at an increased flow rate of 16mL min^-1^ before being returned to the standard flow rate of 3mL min^-1^. Freshly amended soil microcosms were incubated in the dark for a 44-day period under a redox treatment imposed by maintaining headspace gases as needed for each treatment (see treatment section).

#### Treatments and harvests

The impacts of redox periodicity on the soil microbial community and C transformations were examined using four redox treatments: static oxic (SO), low frequency redox fluctuation (LO; 8-days oxic, 4-days anoxic), high frequency redox fluctuation (HF; 4-days oxic, 4-days anoxic), and static anoxic (SA; Fig. 1). ‘Oxic’ conditions were imposed by filling the soil microcosm headspace with ultra-zero air (AI UZ200C), while ‘anoxic’ conditions were imposed using ultra high purity N_2_ (NI UHP250). The introduction of air or N_2_ had previously been used to alter soil redox state to either oxic or anoxic, respectively [2, 21, 26, 39–41].

Soil microcosms were destructively harvested several times throughout the 44-day incubation (red dots on timeline, Fig. 1) in triplicate to capture short-term and long-term temporal dynamics of redox condition. Before removal from flow-through system, microcosms were sealed by capping the jars with air-tight lids to maintain harvest headspace. Microcosms harvested under anoxic conditions were processed in an anaerobic glove box (Coy, Grass Lake, MI), while those harvested under oxic headspace were processed on the benchtop. During harvest, soils were stirred and divided for microbial and chemical analyses. Soils preserved for microbial and metabolite (FTICR-MS) analyses were immediately frozen in the field and stored at −80°C. Iron and phosphorus were sequentially extracted immediately as briefly described below and in detail in companion articles [22, 23].

### Carbon dioxide flux

Ten microcosms were sampled for respiration throughout the incubation experiment. Microcosms were sealed, removed from the flow-through system, and a T_0_ gas sample was taken (30mL) and injected into a pre-evacuated glass vial. Sampled volume was replaced with N_2_ (30mL) to maintain atmospheric pressure in the microcosm. After 1 hour, a second gas sample was collected (30mL), after which microcosms were replaced into the flow-through system. Gas samples were measured on an isotope ratio mass spectrometer at UC, Berkeley.

Carbon dioxide flux measurements were compared across treatments and time by first constraining the data to the desired sample space and then fitting a linear mixed-effects model using the lmer function from the lme4 (v1.1-38) R package. Microcosm was included as a random effect for each model fit and pairwise contrasts were extracted with emmeans (v2.0). All p-values were adjusted with the Benjamini-Hochberg procedure. Cumulative CO_2_ flux was calculated by trapezoidal integration (summing the average flux between each timepoint and multiplying by the time elapsed) and significance was determined using ANOVA and TukeyHSD from the R Stats Package (v4.4.0).

### DNA extraction

DNA was extracted using a modified Griffith’s protocol [42]. Soils from each microcosm were extracted in triplicate then pooled for downstream sequencing. For the extraction procedure, soil (0.25g) was added to lysing matrix E tubes (MP Biomedical, Santa Ana, CA) with 0.5mL 50mM EDTA/350mM KPO_4_/0.7M NaCl/0.5% sarkosyl (w/v), 0.5mL phenol:chloroform:isoamyl alcohol (25:24:1, pH 8), 30µL β-Me, 20µL BSA (400mg mL^-1^). Soils were bead beaten for 45 s at 6.5m s^-1^ in a Fast-Prep (MP Biomedicals, Santa Ana, CA). Sodium chloride (0.7M final) and 100µL 10% CTAB/0.7M NaCl were added to each soil slurry then centrifuged (5 m, 16,000 x g, 4°C). After collecting the aqueous layer, soils were back extracted with NaCl (0.7M) and 0.5mL 50 mM EDTA/350 mM KPO_4_, vortexed and centrifuged. The combined aqueous layers were washed 1:1 with chloroform:isoamyl alcohol (24:1, Sigma Aldrich, St. Louis, MO), emulsified then centrifuged. The collected aqueous layer was mixed with 2 parts 40% PEG 8000/1.6M NaCl and incubated on ice for two hours. Nucleic acids were pelleted by centrifugation for 30 m (16000 x g, 4°C). Pellets were washed with cold 70% EtOH followed by centrifugation twice. Air dried nucleic acid pellets were then resuspended in 50µL TE and replicate extractions were combine (150µL total).

### DNA sequencing

DNA libraries were prepared and sequenced at Argonne National Laboratory (Lemont, IL). Bacterial 16S rRNA gene libraries were prepared using the 515f-806r primer set with a forward barcode [43] and a minor modification to the 806r [44]. Fungal ITS libraries were generated using modified ITS1F and ITS2 primers [45]. Sample libraries were paired end (2 x 250 bp) sequenced on the MiSeq platform (Illumina, San Diego, CA).

### Preprocessing and quality control

Both 16S and ITS amplicons were processed in the same way unless otherwise noted and detailed parameter settings can be found in supplemental methods. Each set of forward and reverse sequences were demultiplexed together using MacQIIME v1.9.1 [46]. Illumina adapters, primers, and PhiX sequences were removed using BBDuk v39.26 (<1% of reads removed from all sequencing runs) [47]. Filtered sequences were run through Cutadapt (v4.6) [48] to remove any primers. Each quality-control filtered set of sequences were further filtered and trimmed using Dada2 (v1.37) [49]. Error rate corrections, dereplication, sample inference, paired read merging, ASV table construction, and chimera removal were performed as outlined in the Dada2 workflow using default parameters for both 16S and ITS samples. Singletons and ASVs found in two or less samples were removed. Bacteria taxonomy was assigned using ‘assignTaxonomy’ from the Dada2 packages with Silva v138 (16S) [50] and fungal taxonomy was assigned using RDP Classifier (v2.14) [51] against the UNITE (v10) (ITS) [52] as the reference taxonomy. A phylogenetic tree was built separately for 16S and ITS amplicons using MAFFT (v7.505) [53] and FastTree (v2.2.0) [54]. ASVs were further clustered at 97% sequence identity using the DECIPHER R package (v3.6.0) [55]. The most abundant ASV within a cluster was selected as a representative, taxonomy and naming scheme was used following clustering, and clustered feature table values are the sum counts of all ASVs within that cluster. The read library size for the 180 samples after quality control ranged from 10,720–208,282 reads per 16S amplicon library (mean of 90,031 reads) and 6,783–175,266 per ITS library (mean of 52,398 reads).

### Bacterial and fungal community analysis

#### Prefiltering

Eukaryotes and Archaea were removed from the clustered bacteria ASV feature table based on Silva taxonomy Kingdom assignment and any ASVs present in two or fewer samples were also removed from further analysis.

Filtered 16S amplicon ASV counts were normalized by variance stabilization transformation using DESeq2 framework [56], and pairwise similarities among samples were calculated using Bray-Curtis dissimilarity. Dissimilarity matrices were visualized using principal coordinate analysis (PCoA). Permutational multivariate analysis of variance (adonis2) [57] statistics were calculated to test the significance of differences among community composition in *a priori* sampling groups. Parameters identified as significant were further tested in pairwise comparisons and the Benjamini-Hochberg (BH) method was used to adjust P-values for multiple comparison correction. Envfit was used to identify individual environmental variables that are significantly correlated with community similarity distance matrices. These analyses were carried out using the Vegan package [57] and Phyloseq package [58] in R v3.4.2. The net related index (NRI) and nearest taxon index (NTI) were measured for the 16S and ITS amplicon data using the Picante package in R [59]. NRI and NTI values were compared across redox treatments and control (T_0_ + Field) samples in the same manner as alpha diversity metrics (richness, Shannon’s Diversity, and Pielou’s Eveness), with ANOVA and either a Tukey HSD (fungal NRI and NTI, bacterial richness) or Kruskal-Wallis Dunn’s test (all other comparisons) depending on data normality and heteroskedasticity.

To assess treatment impacts on discrete taxa abundance the log_2_ fold change (differential abundance) was calculated using DESeq2 compared to the T_0_ pretreatment samples. Specifically, raw, quality-controlled reads were further filtered to keep only taxa present in at least 10% of sample libraries. The log_2_ fold change for each ASV in each treatment/sampling day (compared to the pre-treatment samples) was then calculated.

#### Beta diversity analysis

Filtered ASV tables underwent variance stabilizing transformation via the DESeq2 (v3.19) R package. Briefly, size factors were estimated with estimateSizeFactors type = “poscounts” to account for sparsity. Then homoscedasticity was controlled for with variance stabilizing transformation applied via varianceStabilizingTransformation function (blind = TRUE, fitType = “local”). Bray Curtis dissimilarity was calculated using the vegan (v2.7-2) R package and a pairwise PERMANOVA test was applied via the adonis2 function. R^2^ values, betadispersion metrics, and adjusted p-values (BH and Bonferroni) are in the supplemental (Table S2) PCoA was performed with the wcmdscale function and chemical data were mapped with envfit. Further partial distance-based redundancy analysis of variance on experimental and measured variables (number of days with oxic atmosphere, percent of days with O_2_, pH, DOC, Fe(II), Fe(III), Al, and P) was performed with the dbrda function from the vegan R package (bray_curtis_dist_mat ∼ variable + Condition (day + treatment)).

#### DESeq2 differential abundance analysis

Differential abundance of ASVs in each treatment compared to T_0_ samples was determined via DESeq2. Untransformed feature tables with prefiltering applied were used for this analysis. Prior to DESeq2, feature tables were further filtered to keep only taxa present in at least 10% of samples, size factors were estimated with type = “poscounts” and redox treatment contrasts were performed with fitType = “local” using the Wald test. All plots were generated with ggplot2 (v4.0.1) [60].

ASVs were classified into metabolic strategy categories based on their response patterns in each redox treatment (Fig. S10). Only enrichment data from timepoints where sampling spanned all four redox treatments (days 12, 20, 23, 36, and 44). An ASV was considered enriched if the it had a positive log2 fold change and depleted negative while also having an FDR ≤ 0.1.

#### Iron Utilization Analysis

To determine bacterial ASV’s capacity to use iron, ASV sequences were aligned at 97% sequence identity with at least 80% query coverage to the GTDB 16S rRNA database (ssu_all_r226.fna) [61] using VSEARCH (v2.29.1, -usearch_global, --maxaccepts 5, -- maxrejects 64, --strand both, --top_hits_only) [62]. NCBI accessions from top hits were retrieved using NCBI datasets CLI tool (v18.16.0) [63]. Iron genes were identified in genomes using FeGenie (v1.2) [64] with default parameters. An ASV was considered an Iron Oxidizer if it contained genes in FeGenie categories “iron_oxidation” or “possible_iron_oxidation_and_possible_iron_reduction” with bitscores ≥50. ASVs were considered Iron Reducers if they contained genes in FeGenie categories “iron_reduction” or “possible_iron_oxication_and_possible_iron_reduction” with bitscores ≥50.

The prefiltered ASV feature table was reduced separately to ASVs in either category prior to beta diversity (Bray-Curtis) calculations and ordination (wcmdscale). DESeq2 was run again on the full filtered ASV table (prefiltering and 10% sample occupancy threshold) with contrast groups stratified by sample day and redox treatment. To maximize data retention across these smaller treatment groups DESeq2 was run with minReplicatesForReplace = Inf, independentFiltering = FALSE, and cooksCutoff = FALSE. If an ASV mapped to multiple NCBI genomes, they were included in the iron utilization categories if any of their mapped genomes were identified by the above criteria.

##### Amorphous Fe (AO-Fe) extraction

Poorly crystalline (’amorphous’) iron was extracted from 0.25g soil (dry weight) with 10ml of 0.2M ammonium oxalate solution (pH ∼ 3), shaken for 4 hours, centrifuged at 10,000 rpm for 20 m, and then filtered (0.45μm). Extracts were analyzed for Fe by Inductively Coupled Plasma Atomic Emission Spectroscopy (ICP-AES) at Lawrence Berkeley National Laboratory.

##### Ferrous iron (Fe^2+^) extraction

Fe(II) concentration from this extraction includes soluble and solid-bound Fe(II). Ferrous iron (Fe^2+^) was extracted from 1.0g soil (dry weight) with 5mL of 0.5M HCl, agitated for 30s and incubated in the dark overnight to ensure complete extraction. The supernatant was filtered using a 0.22μm cellulose acetate filter and 100µL or 10µL (when [Fe(II)] >1mmol L^-1^) of the filtrate was added to 0.5mL ferrozine solution (1g L^-1^ ferrozine in 50mmol L^-1^ HEPES buffer, pH 7.0) along with 4mL of MilliQ (18.2Ω) water and incubated for 10 min. Absorbance was measured at a wavelength of 562 nm with a UV-Vis spectrophotometer (Thermo Scientific).

Organic matter characterization: Frozen soils were shipped to EMSL (PNNL, Richland, WA) for metabolite analysis using FTICR-MS as previously described [65]. In brief, soils (300mg) were extracted with water to capture polar metabolites by shaking at 1000 rpm and 23°C for 2 hrs, followed by centrifugation to pellet solids and removal of the supernatant. Water extracts were then diluted with methanol to improve electrospray ionization efficiency. A Suwannee River Fulvic Acid (SRFA) standard (International Humic Substance Society) was injected every 20th sample, followed by a methanol blank, to monitor instrument stability and confirm the absence of sample carryover.

Dissolved organic C in soil extracts was characterized using a 12 Tesla Bruker SolariX FTICR-MS (Billerica, MA). Samples were introduced via an Agilent 1200 series [66] pump through a fused silica capillary (30 μm i.d.) directly into the electrospray ionization (ESI) source at 3.0 μL min-1. Ion accumulation time was optimized per extraction solvent prior to analysis. Instrument parameters were as follows: needle voltage, +4.4 kV to generate negatively charged molecular ions; Q1 set to 50 *m*/*z*; 144 individual transients were co-added for each sample and internally mass re-calibrated using OM homologous series separated by 14 Da (–CH_2_ groups). The mass measurement accuracy was typically within 1 ppm for singly charged ions across a broad *m*/*z* range (*m/z* 200-900).

FTICR-MS peaks were filtered to include those within *m/z* 200-900, those that appear in at least two samples, and to compounds with assigned formulas as determined by ftmsRanalysis (v1.1.0) getVanKrevelenCategoryBounds function with boundary set “bs1”. Water extractible metabolites were annotated based on empirical formula using the compound_calcs function from ftmsRanalysis (v1.1.0) package and compound class differences between treatments were determined via two-way Anova and Tukey’s test. Jaccard distances were calculated from the transformed data and an NMDS ordination was plotted with the vegan R package (v2.7-2) using data transformed to presence/absence (ftmsRanalysis::edata_transform(data_scale = “pres”)). A pairwise permanova was calculated across days and treatments with the pairwiseAdonis (v.0.4.1) package (pairwise.adonis2(jac_wat ∼ treatment * day, data = meta_wat, perm = 999)).

## Results

### Impacts of redox on bacterial and fungal community structure

From the full LEF soil data set (n = 181), we identified 5,248 bacterial and 2,296 fungal ITS ASVs. After clustering ASVs at 97% sequence identity, there were 2,896 and 1,546 bacterial and fungal representative ASVs respectively (Table S1). The bacterial community was dominated by Pseudomonadota (41.3 ± 0.2%), Acidobacteria (18.3 ± 0.3%), Planctomycetes (10.4 ± 0.3%), Verrucomicrobia (9.6 ± 0.1%), Actinobacteria (7.1 ± 0.1%), and Myxococcota (3.4 ± 0%). Ascomycota was the most dominant fungal phylum (41.5 ± 1.1%). Basidiomycota, Mortierellomycota, and Rozellomycota represented 37.3 ±1.4%, 18.5 ±0.7%, and 1.9% ±0.1 of the fungal community, respectively.

Redox conditions significantly impacted beta diversity in both bacterial and fungal communities. Specifically, when considering samples from the full experiment (Field, T_0_ – day 44), SA conditions caused a significant shift in the soil bacterial and fungal community compositions compared to T_0_ and all other redox treatments (FDR < 0.005; Fig. 2, Table S2) and fluctuating treatments did not differ (FDR > 0.05, Fig. 2, Table S2). A similar trend occurred when considering individual timepoints (days 12, 20, 23, 36, and 44) where bacterial communities in the SA condition were significantly different than each other redox treatment (FDR < 0.05, Fig. S1, Table S3). Sampling limitations resulted in low representation of the static treatments (n = 3 SA, n = 3 SO) at day 12, so the PERMANOVA for that day was underpowered and not significant despite clear cluster separation (Fig. S1, Table S3). By day 20 fungal communities in the SA and LF treatments had diverged significantly, and by day 36 the SA fungal communities were different from each other redox treatment (FDR < 0.05, Fig. S1, Table S3). Bacterial communities in the Field condition, HF treatment at day 44, and the LF treatment at day 33 did not differ from T_0_ communities (Figs. S2, S3). However, bacterial communities in each redox condition and from each other timepoint were significantly different from T_0_ communities (Figs S2, S3). Likewise, T_0_ fungal communities were no different than Field condition communities, however they were significantly different from all other redox conditions and sampling days bar SA communities from days 12 and 20 (Figs S1 – S3). We observed soil redox conditions driving changes in microbial community composition in as few as 12 days, and bacterial and fungal communities responded differently to O_2_ availability. Further, prolonged anoxic conditions impacted community composition more than either fluctuating or SO conditions, causing larger community deviations from Field conditions.

**Figure 2.**
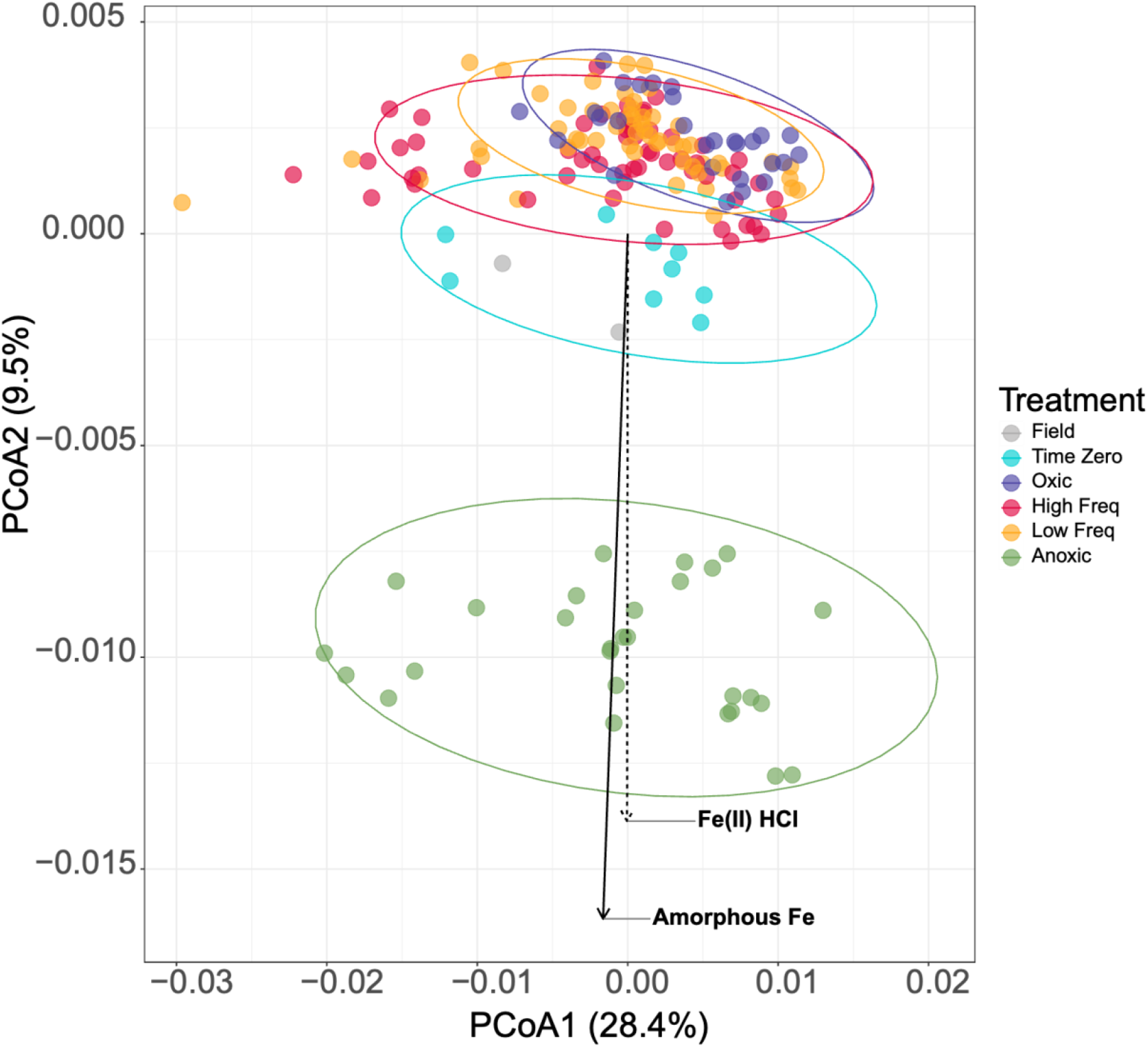
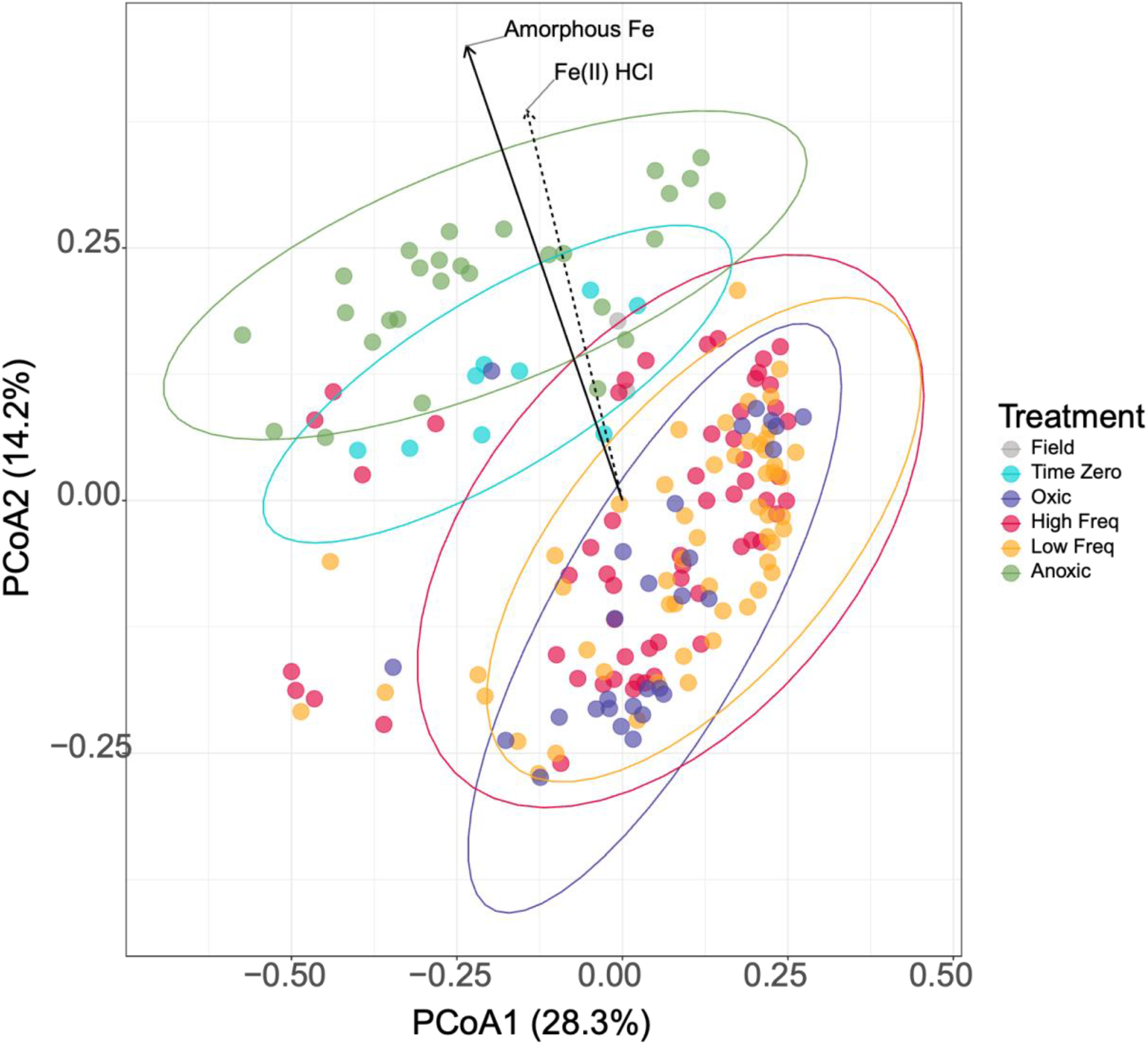
Impacts of redox conditions on the soil microbial community from the Luquillo Experimental Forest in Puerto Rico sampled at days 0, 12, 16, 20, 23, 36, 40, and 44. Field condition soils were sampled after a period of dark incubation for 16days under a four-day oxic and four-day anoxic fluctuating redox regime. After the pretreatment incubation, time all soil was amended with plant litter and Time Zero samples were taken. The remaining 169 microcosms were treated with either static oxic conditions, static anoxic conditions, low frequency (Low Freq) fluctuation between oxic (8 days) and anoxic (4 days) conditions, or high frequency (High Freq) fluctuation between oxic (4 days) and anoxic (4 days) conditions. PCoA of Bray-Curtis dissimilarities calculated from bacterial 16S rRNA gene (top) and fungal ITS gene (bottom) counts. To determine the influence of the measured environmental chemistry on community composition, we calculated ordination loadings for all metabolite classes, number of days with oxic atmosphere, percent of days with O_2_, pH, DOC, Fe(II), Fe(III), Al, and P. The plot shows Fe(II) and Fe(III) (P = 0.001, R^2^ = 0.67 and 0.81, respectively) for the bacterial communities (top) and the fungal communities (bottom) Fe(II) and Fe(III) (P = 0.001, R^2^ = 0.29 and 0.38, respectively). Ellipses are multivariate t-distribution of each treatment at 0.95 confidence.

We assessed the beta-dispersion of the microbial communities within each treatment to determine how cohesive community responses were between treatments (Fig. 2, Table S2). The dispersion between treatments was significant for the SA bacterial community compared to the T_0_ and SO communities, and there were no significant differences in dispersion in the fungal communities. Overall, bacterial and fungal community composition were influenced by soil redox conditions and the most pronounced shifts in bacterial communities occurred in the SA treatment, though differences between SA and either SO or T_0_ may be influenced here by intra-group variance (Fig. 2, Table S2).

We also tested several experimental and environmental variables (number of days with oxic atmosphere, percent of days with O_2_, pH, DOC, Fe(II), Fe(III), Al, and P) for their impact on the overall observed bacterial and fungal community structure. Of these, the percent of days with O_2_ and both species of iron had the strongest correlation to bacterial community ordination clustering, and fungal community ordinations correlated strongest with number of days with oxic atmosphere, percent of days with O_2_, and iron (Envfit, FDR < 0.01, Additional Table 1). Of the environmental measurements taken, Fe(II) and Fe(III) accounted for the most variation in both bacterial (R^2^ 0.67 and 0.81, respectively) and fungal (R^2^ 0.29 and 0.38, respectively) communities (Fig. 2, Additional Table 1). To assess the impact of redox treatment and timepoint on the correlation of these variables to the microbial communities, we performed a partial distance-based redundancy analysis. Once variance associated with the conditional variables (day and redox treatment) were accounted for, the experimental and environmental factors previously associated with community structure no longer correlated significantly to either bacterial or fungal communities (Additional Table 1). Soil iron and time exposed to oxic atmosphere covary strongly with redox treatment and the primary drivers of community variation may instead be the redox conditions and the duration spent in each condition.

Bacterial community richness and evenness did not vary significantly from the Field condition or T_0_ (Fig. S4A). However, Shannon’s Diversity was significantly higher in the SO treatment compared to control samples (Fig. S4A). In pairwise comparisons between treatments, community richness was significantly higher in the SO condition compared to HF and LF (ANOVA-Tukey, *P* < 0.05). Community evenness was higher in HF compared to SO (Dunn’s test, P < 0.01) and in LF compared to SO (Dunn’s test, P < 0.01). The fungal communities did not vary in either richness, Shannon’s Diversity, or evenness across treatments (Fig. S4B). SO conditions supported more taxa in bacterial communities relative to changing conditions, but redox fluctuation allowed some bacteria to occupy a higher proportion of the community relative to clades that dominate periods of consistent O_2_ highlighting community resilience to fluctuating redox.

Redox impacted phylogenetic community structure in both fungal and bacterial communities. SA bacterial communities exhibited higher NTI and NRI compared to the other redox treatments (Dunn’s test, P < 0.05, Fig. S5A), and control communities had higher NTI than fluctuating and SO communities (Dunn’s test, P < 0.01, Fig. S5A). The high NRI and NTI values in SA soils compared to other treatments suggest prolonged anoxic conditions are a selective pressure favoring certain bacterial lineages, and that even intermittent periods with O_2_ can mitigate this effect. The fungal communities displayed no NRI or NTI differences between any treatments (ANOVA-Tukey, P < 0.05, Fig. S5B). This suggests LEF soil fungal communities are more resilient to extended periods of anoxia than bacterial communities are.

### Taxon specific response to redox

Using differential abundance analysis, we identified bacterial and fungal taxa that were either significantly enriched, depleted, or unchanged for each redox treatment compared to T_0_ (Fig. 3 & Fig. S6, respectively). The majority of fungal and bacterial ASVs observed were not impacted by redox treatment (Fig. 3, Fig. S6). Specifically, a total of 199 bacterial ASVs (spanning 16 phyla) and 56 fungal ASVs (spanning six phyla) were significantly impacted (FDR < 0.05, |log2FoldChange| ≥ 2). Relative to T_0_ conditions, we observed more bacterial ASVs being significantly enriched (FDR < 0.05, log2FoldChange ≥ 2) in the SA treatment (112 ASVs) compared to either SO (80 ASVs), HF (81 ASVs), or LF (81 ASVs) redox treatments. Interestingly, the opposite trend was observed with in the fungal communities where the most ASVs were enriched in LF (34 ASVs), followed by HF (23 ASVs) and SO (23 ASVs), and finally SA (2 ASVs). Only a single bacterial ASV from the SA condition was significantly depleted (FDR < 0.05, log2FoldChange ≤ −2) compared to T_0_. Conversely, 24 fungal ASVs were significantly depleted: 9 in SA, 6 in LF, 5 in HF, and 4 in SO. We observed bacteria and fungi in these soils to be most sensitive to SA conditions compared to the other redox states. Within the community membership most sensitive to redox, differences in SA fungal community composition were driven by depletion of taxa rather than enrichment and the opposite is true for bacterial communities.

**Figure 3.**
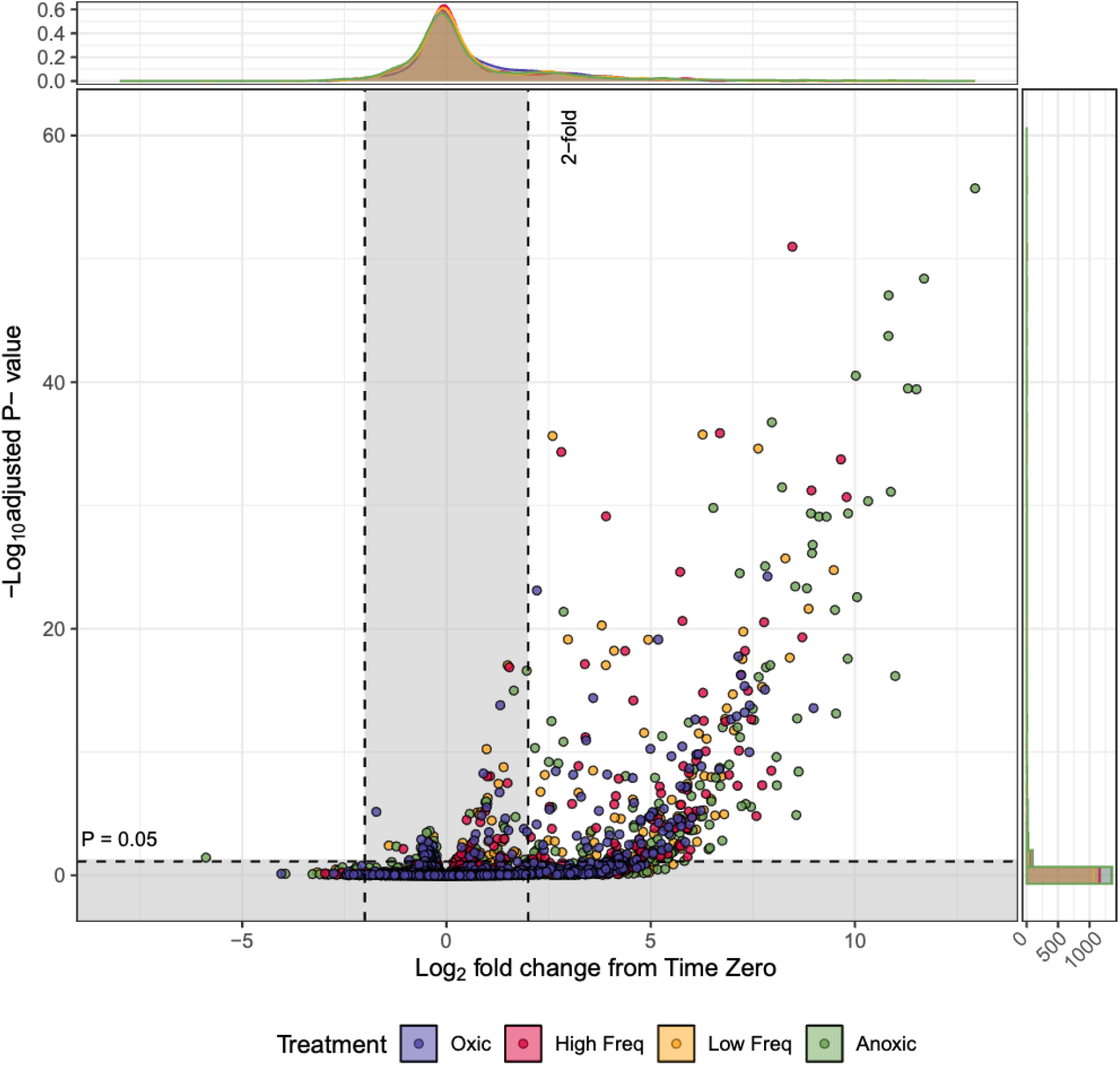
Significant (unshaded plot area) and nonsignificant (shaded area) Log_2_ fold change response of bacterial ASVs in each treatment (colors) relative to T_0_ identified using DESeq2. P-values were adjusted for multiple comparisons using ‘Benjamini-Hochberg’ correction.

Looking at only the ASVs that were significantly enriched or depleted (Fig. 3 and Fig. S6), we identified the specific taxa significantly impacted by each redox treatment throughout the experiment (Fig. 4). Of the 16 phyla and unclassified ASVs significantly impacted by redox, Bacillota, Chloroflexota, Thermodesulfobacteriota, Actinomycetota, and Fibrobacterota ASVs were solely enriched in the SA treatment, and the only depleted ASV was from Bdellovibrionota. Notably, of the 57 ASVs enriched only in the SA treatment, 51 were Bacillota. Membership from the Chlamydiota phyla were only enriched in SO conditions. The fungal phylum Ascomycota represented more than half (n = 33) of all fungal ASVs significantly impacted by redox (n = 60), and 26 of those were significantly enriched in at least one redox condition compared to T_0_ condition. Of those, a disproportionate number of Ascomycota ASVs were enriched in treatments with O_2_ (LF n = 20, SO n = 14, HF n = 13, SA n = 1). Interestingly, Glomeromycota ASVs were depleted in each treatment bar HF (LF n = 1, SO n = 3, SA n = 4) and enriched in no treatment compared to T_0_. In any condition with even some O_2_ (HF, LF, SO), at least one member from Ascomycota, Mortierellomycota, Basidiomycota, and Incertae Sedis was significantly enriched (Fig. 4B).

**Figure 4.**
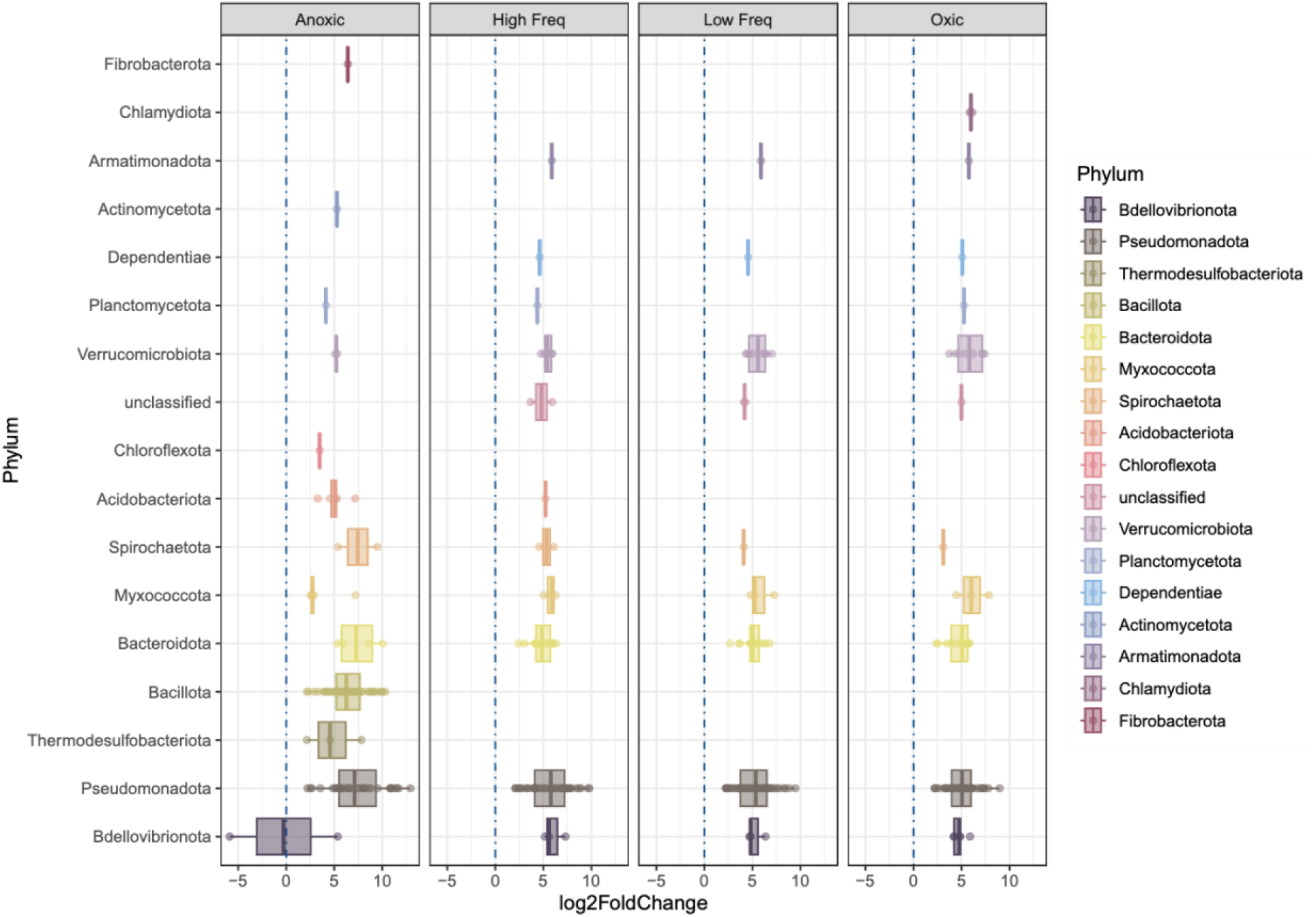
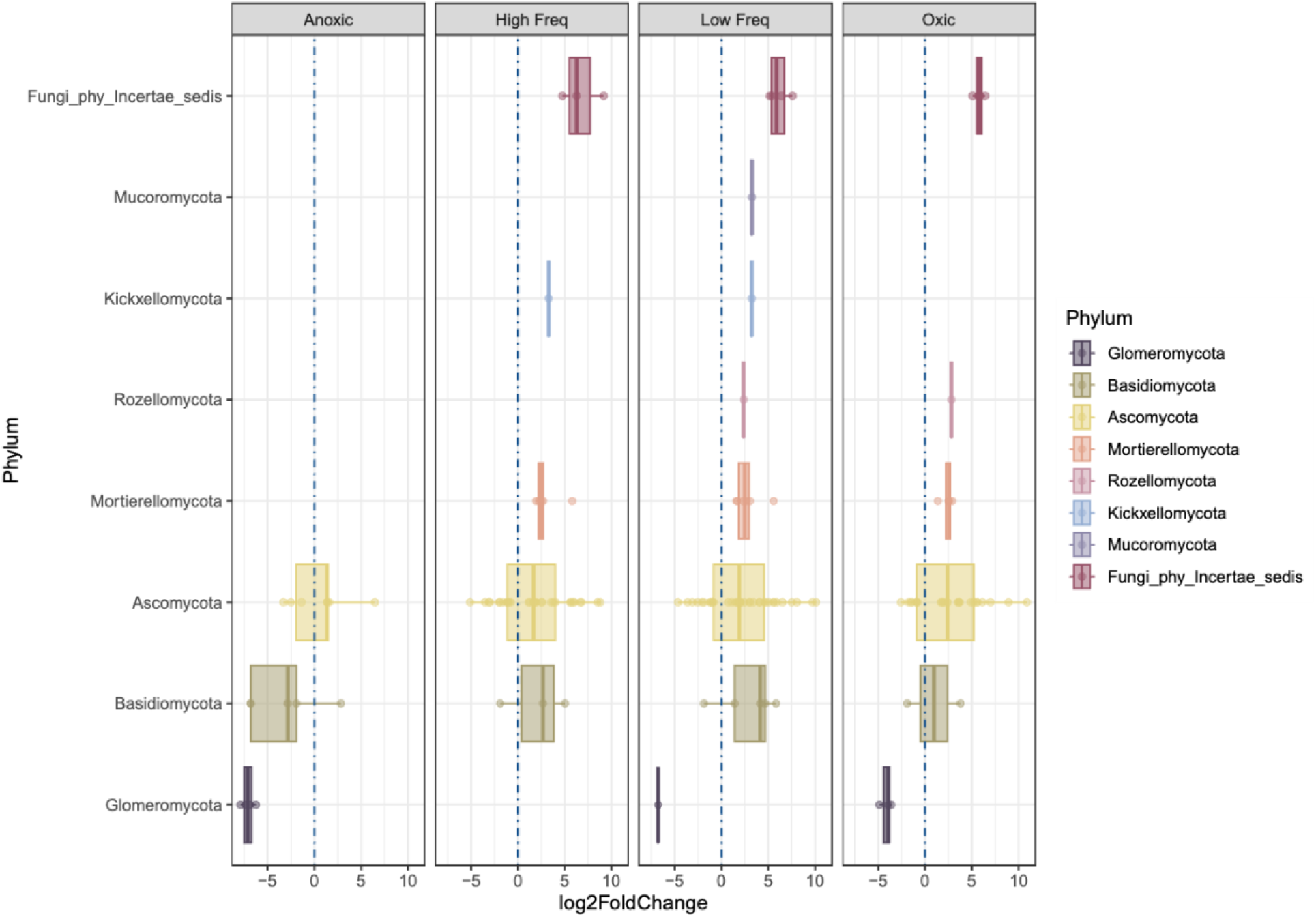
Bacterial (A) and fungal (B) ASVs identified using DESeq2 as significantly enriched (right of blue dashed line) or depleted (left of blue line) relative to their abundance at T_0_. Points represent actual log2 fold change value and box plots represent interquartile range of log2 fold change for significant ASVs in each taxonomic order.

We classified ASVs with significant enrichment (per differential abundance analysis) into metabolic strategy categories based on their response patterns in each redox treatment (Table S4, Fig. 5). Microbes were considered “Plastic” if they were enriched in both SO and SA treatment and either, neither, or both fluctuating treatments. “Facultative Advantage” microbes were classified if they were enriched in either or both fluctuating treatments and not the static treatments. “Obligate Anaerobes” were only enriched in the SA treatment, and “Obligate Aerobes” were only enriched in the SO treatment. “Tolerant Aerobes” were enriched in the SO treatment and at least one of the fluctuating treatments but not enriched in the SA treatment, and “Tolerant Anaerobes” were enriched in the SA treatment and at least one of the fluctuating treatments but not enriched in the SO treatment. Response patterns were assessed for each day of the experiment, and ASVs were counted if they were significantly enriched or depleted (FDR < 0.1, log2 FC compared to T_0_). Tolerant Aerobe (n=144 ASVs) was the highest represented response pattern throughout the experiment followed by Facultative Advantage (n=126 ASVs). Thermodesulfobacteriota were the only clade with representation solely as Obligate Anaerobes. Bacillota spp. were highly represented as Obligate Anaerobes, and Pseudomonadota was the clade with the most ASVs classified in every category bar Obligate Anaerobe. Chloroflexota ASVs tended towards anoxic conditions adopting Obligate Anaerobe, Tolerant Anaerobe, and Facultative Advantage metabolic strategies. Of the fungal ASVs classified into metabolic strategy categories, the phyla with the most ASVs in each category was Ascomycota. Mortierellomycota were mostly Facultative Advantage and Tolerant Aerobe while Basidomycota were mainly Facultative Advantage and Obligate Anaerobes. These groupings allowed us to determine that most fungal and bacterial membership can thrive in soils with oxic and anoxic periods and that static periods of high or low redox enriches a subset of the broader community.

**Figure 5.**
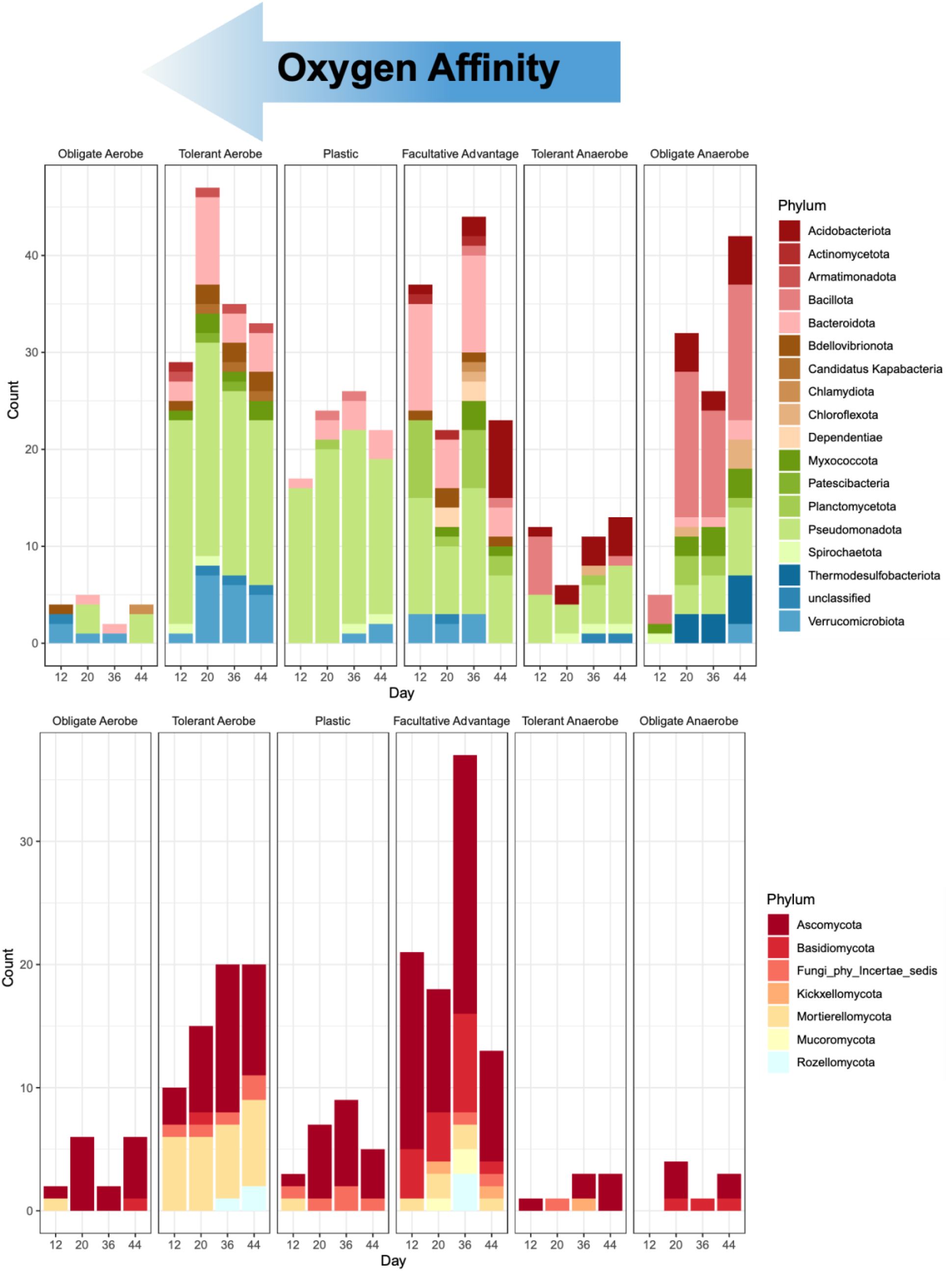
Number of bacterial (top) and fungal (bottom) ASVs in each metabolic strategy category.

### Impacts on Fe reduction and oxidation

To resolve the biological drivers behind the observed Fe-redox relationship, we examined the changes of inferred Fe-cycling bacteria for each of the treatments across days (Fig. 6). Community ordinations of bacteria linked to genomes with either Fe-oxidizing or Fe-reducing genes show the SA treatment significantly differs from both T_0_ and other redox conditions (FDR < 0.05). In all treatments there was a net decrease in Fe-oxidizers and all but SO showed a slight increase in Fe-reducers (Table S5). In SA conditions Fe-reducers experienced a net increase of 19 ASVs relative to T_0_ (Fig. 6, Table S5). Fe-reducing bacteria enriched in SA hail from Bacillota (n=9), Acidobacteriota (n=3), Myxococcota (n=3), Thermodesulfobacteriota (n=4), Pseudomonadota (n=3), and Verrucomicrobiota (n=1). SO conditions yielded fewer Fe-reducing and Fe-oxidizing bacteria, one Acidobacteriota Fe-reducer ASV was depleted in SO soils and six Fe-oxidizer ASVs from Acidobacteriota (n=4), Latescibacterota (n=1), and Verrucomicrobiota (n=1) were depleted in SO soils. It may be the case that Fe-oxidizing bacteria are depleted in the fluctuating and SO treatments due to rapid abiotic Fe oxidation when O_2_ is present [22]. Conversely, anoxic conditions both enrich for Fe-reducing bacteria and deplete Fe-oxidizing taxa suggesting a potentially important role for Fe as a terminal electron acceptor for anerobic respiration possibly allowing bacteria able to utilize oxidized Fe in this way to outcompete taxa dependent on O_2_ to respire.

**Figure 6.**
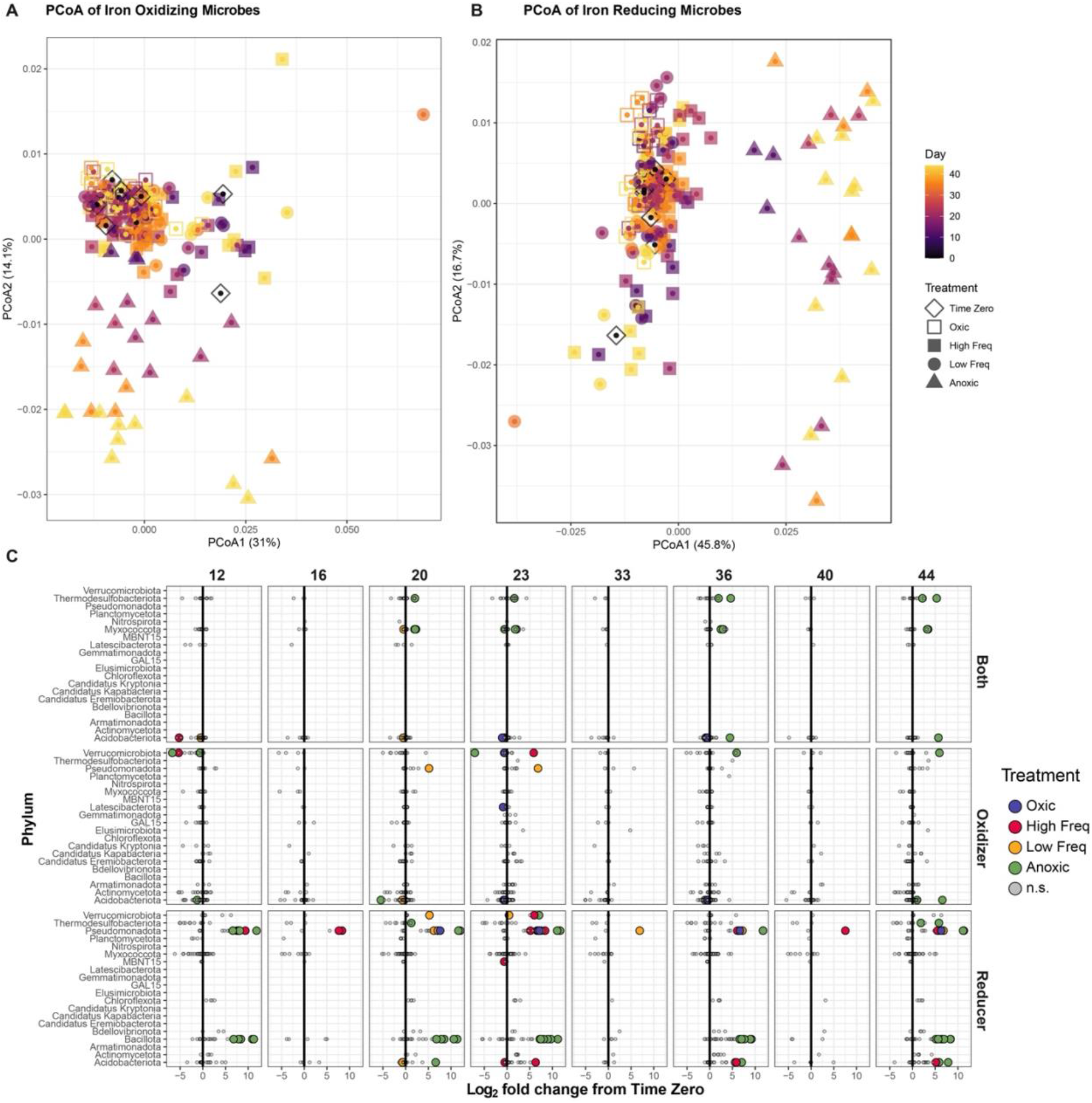
Impacts of redox conditions on (A) iron oxidizing and (B) iron reducing bacteria identified by 16S rRNA genes in a wet tropical soil. Soils were incubated for 44 days under four redox treatments: two static conditions (oxic, anoxic) and two redox fluctuating conditions, with either high frequency or low frequency oscillation. (C) Specific responses of iron oxidizing and reducing taxa over time (incubation day is noted at the top of each figure); colored by the treatment when it is significantly enriched (to the right of bold black line) or depleted (to the left of the line) relative to Control, non-significant (n.s.) ASVs are colored gray.

### Dynamics of organic matter cycling: respiration and metabolite transformations

Continuous gas measurements revealed distinct temporal mineralization dynamics across redox treatments (Figs. 7A, S7A). Soil CO_2_ fluxes decreased over time, regardless of O_2_ headspace conditions. The SO and two fluctuating treatments exhibited exponential decay in CO_2_ respiration rate stabilizing around day 20. Our modeling indicated that redox treatment (P = 0.009) and time (P < 0.0001) significantly impacted CO_2_ fluxes and exhibited a significant interaction (P < 0.0001). Overall, CO_2_ flux in the SA treatment were 0.065 μmol C g^-1^soil hr^-1^ (SE = 0.021, p = 0.006) higher than in the HF and 0.073 μmol C g^-1^soil hr^-1^ (SE = 0.02, p = 0.003) higher than in the LF treatments and the SO treatment was 0.046 μmol C g^-1^soil hr^-1^ (SE = 0.021, p = 0.044) higher than in the HF treatment and 0.054 μmol C g^-1^soil hr^-1^ (SE = 0.021, p = 0.02) higher than in the LF treatment. When all time points are included in the model, SA and SO treatments show no significant difference in CO_2_ fluxes. Considering only later timepoints (day 20 – 44), SA CO_2_ fluxes were 0.058 μmol C g^-1^soil hr^-1^ (SE = 0.017, p = 0.008) lower than SO. Modeling the CO_2_ flux across days in each redox treatment showed that SA conditions had 0.096 μmol C g^-1^soil hr^-1^ (SE = 0.036, p = 0.025) lower CO_2_ flux compared to SO at day 3, while the fluctuating treatments did not differ from SO at that timepoint (Fig. S7B). As the experiment progressed to day 12, SA had significantly higher CO_2_ emissions than SO (0.051 C g^-1^soil hr^-1^ ±0.023, p = 0.042), HF was no different from SO, and CO_2_ fluxes in LF were 0.059 C g^-1^soil hr^-1^ (SE = 0.024, p = 0.042) lower than in the SO treatment. By day 24 SA CO_2_ did not differ from SO, but CO_2_ fluxes in both fluctuating treatments were significantly lower compared to SO. As time progressed, SA CO_2_ emissions continued to decline significantly compared to SO, while the fluctuating treatments and SO CO_2_ levels became significantly indistinguishable. When we considered the cumulative CO_2_ emissions from each redox treatment, the only significant difference was observed between SO and LF treatments (Fig. 7B). Early suppression of CO_2_ production under SA conditions may reflect rapid O_2_ depletion and a shift away from aerobic metabolism. Further, the relative decline in community respiration rate at midpoints (day 12 and 24) in fluctuating treatments compared SO may reflect a shift towards facultative anaerobic or fermentative pathways. The convergence of fluctuating and SO respiration levels in later timepoints suggests establishment of a new metabolic steady state, whereas the continued decline in the SA levels indicate prolonged anoxia exerts sustained constraints on community respiration. Conversely, the near indistinguishable cumulative CO_2_ differences across treatments may point to a resilient microbial cohort able to sustain activity despite redox variance.

**Figure 7.**
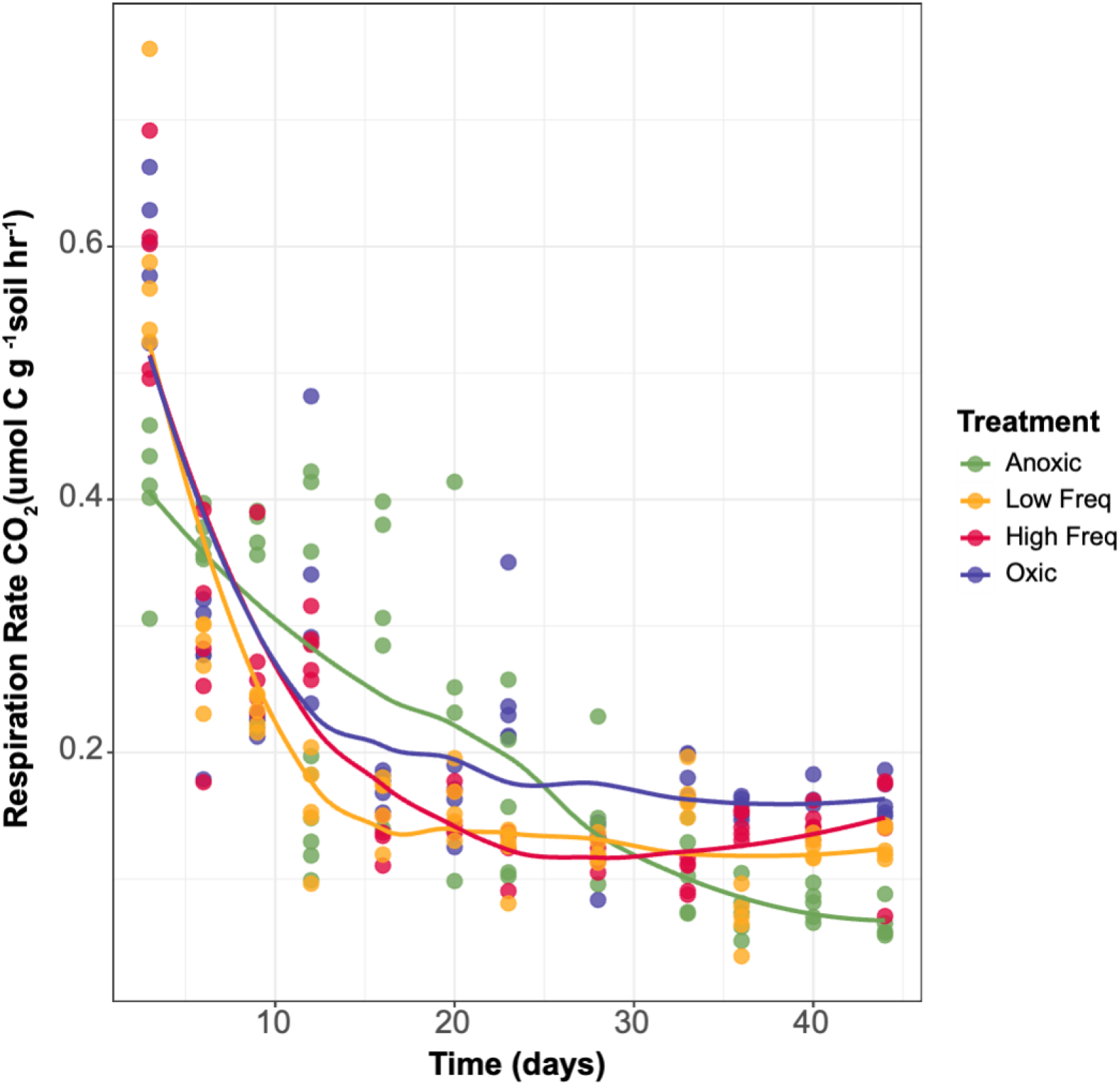
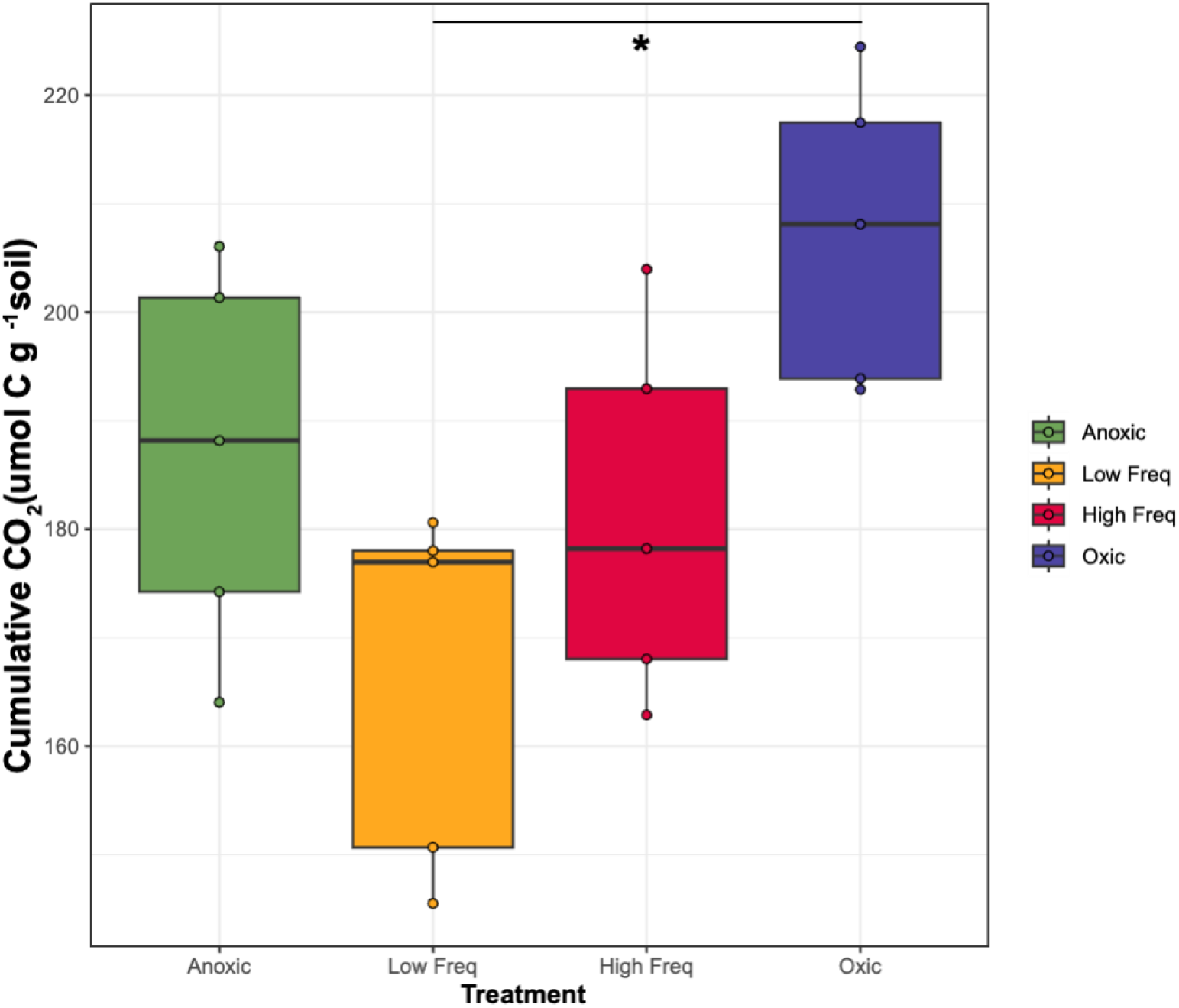
Ten microcosms were sampled for respiration throughout the incubation experiment. A) CO_2_ measurements over 44-day experimental period were sampled and compared across treatments and time. B) Cumulative CO_2_ flux was calculated by summing the average flux between two time points and multiplying by the time elapsed, * p < 0.01

FTICR-MS-based organic matter characterization revealed that redox conditions shaped soil organic matter composition (Fig. 8). A pairwise PERMANOVA comparing Jaccard distances between organic matter profiles from different treatments and timepoints showed redox significantly impacted organic matter composition between treatments and SA soils exhibited the strongest divergence from each of the other conditions (R^2^ = 0.55 – 0.59, P = 0.012 - 0.001). A direct comparison of organic matter compound classes between treatments indicated that the SA treatment had significantly less unsaturated hydrocarbon–like compounds than both fluctuating treatments and the SO treatment (Fig. 8, P < 0.001). The SA soils had significantly higher levels of protein- and carbohydrate–like compounds than each of the other redox treatments and T_0_ (Fig. 8, P < 2e-16), significantly higher tannin-like levels than HF and T_0_ soils (Fig. 8, P = 0.01 – 0.007), and significantly higher levels of lignin- and amino sugar-like compounds than each of the other redox treatments (Fig. 8, P < 0.001). The relatively high abundances of certain compounds like amino sugar-, carbohydrate-, tannin- and lignin-like compounds in the SA condition may be indicative of reduced microbial activity or a diminished enzymatic potential to break down certain C sources (i.e. less laccase activity breaking down lignin because of reduced O_2_)[67]. Conversely, higher levels of compounds like unsaturated hydrocarbon-and lipid- like in soils with O_2_ could indicate higher cellular turnover in those treatments.

**Figure 8.**
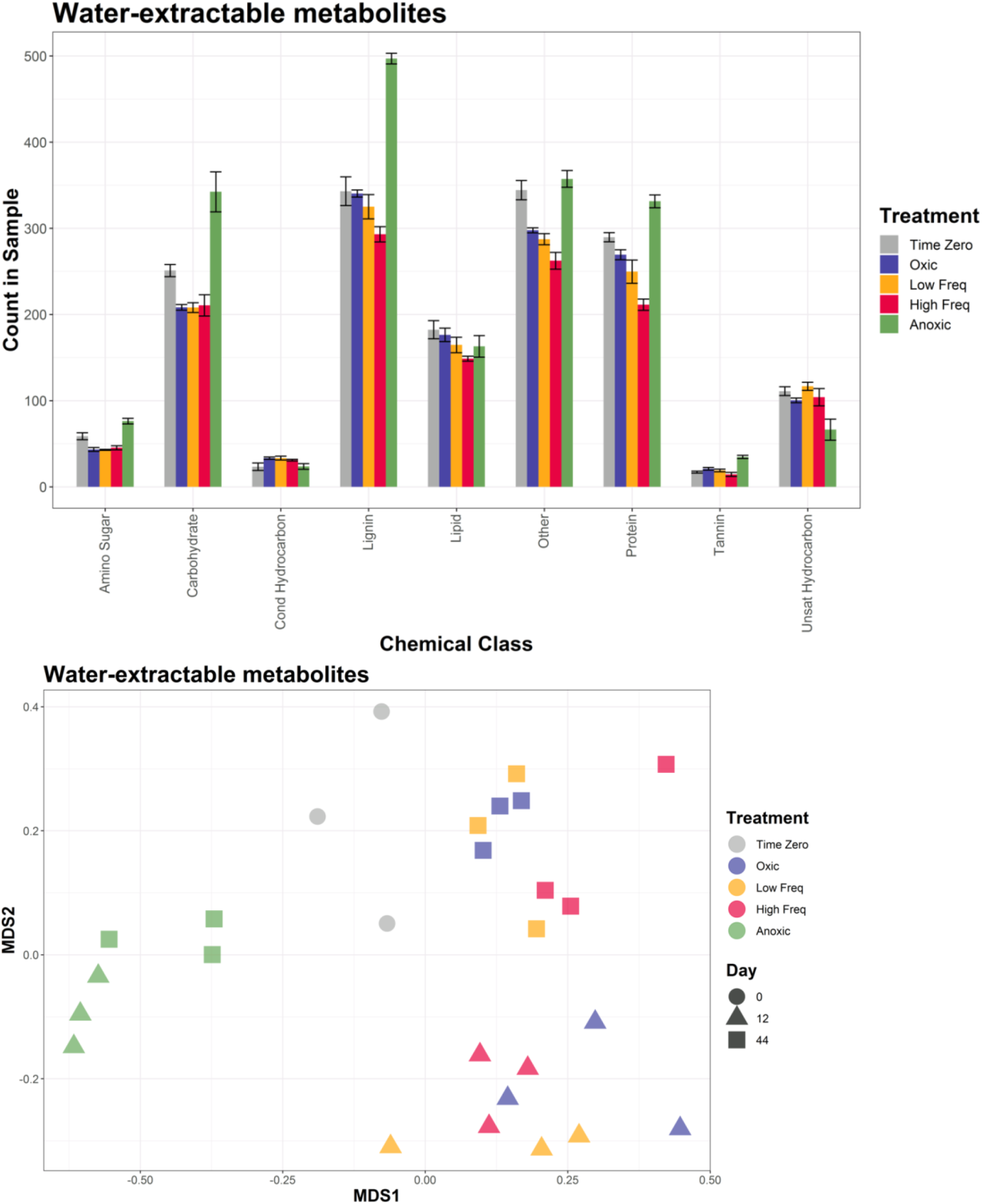
Impacts of redox conditions on the water extractable soil metabolites composition from the beginning of the experiment (T_0_ and day 12) to the end (day 44) were determined by FTICR-MS and peak classification into compound class-like species using Van-Krevlin space. Bars indicate the number of compounds identified within each compound class from days 0 and 44 (top). NMDS of presence/absence-normalized water extracted metabolites matrix (stress = 8.48e-5) from days 0, 12, and 44 (bottom).

## Discussion

Climate modeling predicts that as global maximum temperatures increase, tropical soil C stocks will likely decrease [68]. This represents a transfer of organic C from one of the largest global repositories, tropical soils, to greenhouse gases like CO_2_ and CH_4_ (methane) mediated by microbial activity [69–71], highlighting the importance of soil microbial community structure and function to broader biogeochemical processes. Abiotic factors other than temperature that contribute to soil microbial community structure and C use include pH, soil moisture, and substrate type [72, 73]. Conveniently, soil redox potential represents an environmental condition directly influenced by pH and soil moisture (via O_2_ availability), that dictates organic and inorganic substrate utilization by microbial metabolisms contributing to elemental cycling. Here we showed that microbial communities in LEF soil change with static redox conditions in a manner that correlates with redox-sensitive processes involving C and N transformations [21]. As such, it is important to understand how various redox conditions impact the tropical soil microbiome and how soil microbial community structure adapts to different redox environments, enriching membership with specific metabolisms that can dictate the fate of soil organic C and influence biogeochemical cycles.

Our previous work has demonstrated the soil microbial community present in LEF soils is stable under dynamic redox conditions [2, 21]. Our data support these previous findings and suggest that many members of this microbial community thrive despite frequent redox variability, as evidenced by the larger number of fungal taxa enriched in the fluctuating redox treatments compared to the static redox treatments, and our observation that most fungi and bacteria we detected were not impacted by redox or occupied facultative, plastic or tolerant ecological categories (Figs. 3, 5, S9). Conversely, there are subsets of the community that are more adapted for SO or SA conditions. While Bacillota and Pseudomonadota were enriched in the SA soil, the SO conditions notably enriched fungi in the Hysteriales and Mortierellales orders and bacteria in the Chlamydiales order (Fig. 4). Hysteriales are part of the Ascomycota phylum, one of the most phylogenetically diverse fungal groups and broadly considered a more drought-tolerant phyla [74, 75]. Our data indicate saprotrophic Helotiales and Xylariales of the Ascomycota phylum were depleted in both static redox conditions, highlighting the diverse life strategies and redox sensitivities within this clade. Interestingly, a study conducted across 43 sites from the Qinghai-Tibetan Plateau wetland revealed that increased C:N ratio favored Hysteriales spp [76]. As such, Hysteriales in LEF soils may gain a competitive advantage as N is depleted by microbial metabolic activity during SO conditions, increasing the C:N ratio.

In the field, increased soil O_2_ occurs with soil drying resulting from lower precipitation input in addition to soil drainage. The SO and fluctuating communities in our study did not differ significantly from the native soil community and T_0_ (Fig. 2), suggesting they were more resilient to periods of high redox (i.e., high O_2_ availability). Further, our data show specific bacterial and fungal clades enriched in treatments with O_2_, such as Armatimonadota, Dependentiae, Mucoromycota, Kickxellomycota, Rozellomycota, Mortierellomycota, and several unclassified bacteria and fungi (Fig. 4). Similarly, the anoxic communities enriched bacteria like Bacillota, Thermodesulfobacteriota, Chloroflexota, and Fibrobacterota (Fig. 4). In a rainfall exclusion experiment in the LEF, a ‘drought’ response was measured for distinct microbial groups, but not in the overall microbial community [77]. A study that imposed reduced precipitation in forests across three regions of Germany (northeast, central, and southwest) for six months found negligible impacts on the total bacterial community while there were significant impacts on a subset of the active bacterial community [20]. Our data substantiate these previous findings and although soil moisture was not manipulated in this study, soil moisture, soil O_2_, and soil redox are not necessarily independent of one another [17, 72]. LEF soil microbial communities appear resilient to changes in soil redox like those resulting from moderate to severe drought and redox impacts seem limited to a susceptible subset of community membership like those enriched in either static condition.

Our data indicate sustained anoxia acts as a strong environmental filter in these soils, enriching for bacterial strains in the Bacillota and Pseudomonadota and depleting saprotrophic fungi from Ascomycota. Additionally, of the environmental variables we measured, Fe(II) was correlated most strongly with the SA community structure shift (Fig. 2). This coincided with the enrichment of Fe-reducing microbes (Klebsiella spp., Geobacter spp., and Desulfovibrio spp.) and certain Fe-oxidizing microbes (Chromobacterium spp., Dechloromonas spp., and Rhodomicrobium spp.) in the anoxic treatment during later timepoints (Days 12-44, Fig. 6C) suggesting iron could play an important role for microbes during anoxic periods. Overall, the iron reducing community in the anoxic treatment was very different from the other treatments by the end of the experiment, while the change observed in iron oxidizing bacteria was less pronounced across treatments (Fig. 6B). These results indicate a potential shift in iron utilization by the community during extended periods of anaerobic conditions, supporting iron as a likely electron acceptor allowing enrichment of iron reducing bacteria. The correlation of anoxic community structure with Fe(II) supports this supposition, as an accumulation of iron in its reduced valence state could occur as iron reducing bacteria respire Fe(III) (ferric iron) causing Fe(II) accumulation sans abiotic oxidation and reduced microbial mediated iron oxidation during the extended soil anoxia.

A study conducted on soils from rice paddies highlighted that of the seven anaerobic or facultative anaerobes and of the four chemoautotrophic bacteria that were enriched following Fe(II) addition, four and three respectively were members of Pseudomonadota (formerly Proteobacteria) or Bacillota (formerly Firmicutes) [78]. This notion was mirrored in a separate study of tundra soil microbiomes that noted higher relative abundances of Bacillota and anaerobic iron reducing Deltaproteobacteria in persistently anoxic tundra [79]. Although the anaerobic metabolic process of dissimilatory iron reduction has primarily been studied in submerged sediments, it has been shown to sustain high rates of microbial respiration in surface horizons of terrestrial humid soils, where densities of iron-reducing bacteria can exceed those of submerged sediments [27, 33, 40]. Rates of CO_2_ production from iron reduction can be highly significant: anaerobic respiration rates from a tropical Ultisol measured 70–100% of aerobic controls over short-term incubations [80]. Further, short-range order iron oxides are often abundant in humid tropical soils and can eventually be depleted by reduction and leaching [81], potentially limiting the overall importance of iron reduction and contribution to soil C accumulation [4]. Concentrations of short-range order minerals declined with increasing precipitation along the same gradient, implying that anaerobic constraints on decomposition as opposed to organo-mineral interactions were primarily responsible for increased carbon accumulation [4]. These findings together with the community structure shifts we observed after prolonged anoxia in LEF soils may indicate that the ability of membership to utilize plant litter effectively as a C source in O_2_ limited conditions is linked to their efficient use of the various oxidation states of iron, either iron(II) or iron(III) as reductant or oxidant respectively. Initial competition for oxygen as an electron donor for ferrous chemoautotrophy may select early for iron oxidizing bacteria before O_2_ is fully depleted, while success later during anoxia may hinge on the ability to then reduce ferric iron [82].

Our FTICR-MS-based organic matter characterization of water-extractable compounds revealed a distinct soil organic matter composition in anoxic soils as compared to the other redox treatments, demonstrating that O_2_ availability is a key determining factor in soil C transformations. Soil C is a highly heterogenous pool of different C compounds and only a very small fraction of this C is readily accessible for microorganisms [30, 83]. Lignin, for instance, is a phenolic component of lignocellulouse that is particularly challenging for microbial metabolism and often lags behind total liter decomposition rates [84]. Bacteria able to utilize lignin initiate lignin depolymerization with oxidases or peroxidases like laccase or lignin peroxidase [85]. These enzyme classes are predicated on O_2_ availability either directly as an electron sink during substrate oxidation or indirectly during the production of hydrogen peroxide which then facilitates substrate oxidation as the terminal electron acceptor of peroxidase [86]. We observed SA soils containing higher levels of amino sugar-, carbohydrate-, lignin-, protein- and tannin-like compounds than both SO and fluctuating treatments. The elevated amino sugar pools in the SA water-extractable fraction likely reflect reduced decomposition of microbial cell wall material under anoxic conditions, suggesting that O_2_ limitation constrains the turnover of not only plant-derived but also microbial-derived organic matter.

Also, anoxic soil CO_2_ respiration rate decreased more slowly than the precipitous rate drops in either oxic or fluctuating soils as time passed (Fig. 7). These data may reflect the rate-limiting oxidative depolymerization of lignin and other complex carbohydrates and recalcitrant aromatic compounds, where similar O_2_ dependent metabolic enzymes not only account for higher lignin and other C substrate levels in our anoxic soils, but also for the lagging decline in C mineralization compared to O_2_ containing treatments (Fig. 7). Furthermore, the reductive dissolution of Fe(III) minerals under anoxic conditions likely released previously mineral-associated organic matter back into the water-extractable pool, contributing to the elevated carbohydrate and protein levels we observe. This represents a direct mechanistic link between the iron reduction dynamics documented in our microbial community data and the organic matter composition shifts detected by FTICR-MS.

This offers a mechanistic explanation for slower litter decomposition during O_2_ limited periods, the higher DOC levels in the SA treatment, and the higher CO_2_ flux during SO periods. Conversely, the higher levels of unsaturated hydrocarbon- and lipid-like compounds in the water-extractable fraction of SO soils point toward active microbial membrane biosynthesis and biomass turnover under aerobic conditions, consistent with the elevated CO_2_ flux and greater community respiration rates observed in those treatments. Slower C turnover in the anoxic treatment may also be a bellwether to microbial membership shifts to anaerobic constituents with cellular machinery more adapted to utilize alternative electron acceptors for respiration as well as C substrate metabolism.

## Conclusion

Here we manipulated the redox conditions in tropical soils for 44 days and examined changes in C turnover, microbial community structure, and soil chemistry to determine the impact of O_2_ availability on microbially mediated biogeochemistry. LEF soil microbial communities are resilient to periodic redox shifts; though extended anoxic periods significantly changed microbial community composition. We observed specific clades respond to both SA and SO conditions. Soil redox patterns strongly impacted C cycling, liberating more CO_2_ when oxic, and more DOC when anoxic. Redox status determined both the quantity and composition of water-extractable organic matter, with iron reduction under anoxic conditions releasing mineral-associated compounds into the dissolved pool and O_2_ availability governing the enzymatic capacity to decompose recalcitrant C. Our findings highlight the importance of soil redox conditions to microbial community structure, function, and influence on C mineralization and SOM composition. More broadly, our results suggest that microbial adaptation to fluctuating redox environments plays a critical role in determining how C is processed and stored in tropical soils, linking community assembly to ecosystem biogeochemical function. As rainfall patterns in tropical zones increase in intensity and frequency, so too will extended periods of anoxic soil conditions. Likewise, if we continue to observe extended drought periods these prolonged phases of either anoxic or oxic soil conditions will create circumstances that we have shown disrupt the soil microbial community away from the established cohort that is robust to periodic redox fluctuation. Further research is warranted to identify the specific mechanisms most impacted by soil redox and to model C fate in the face of a changing climate. Understanding these ecological responses will be critical for predicting how tropical soil C storage and greenhouse gas fluxes respond to future climate change.

## Acknowledgements

This study was supported by an Early Career Research Program award to JPR from the U.S. Department of Energy, Office of Biological and Environmental Research (DOE – OBER). Additional support for manuscript preparation was provided by the DOE OBER Genomic Science Program “Microbes Persist” Scientific Focus Area (#SCW1632) at LLNL. A portion of this research was performed on a project award (48643) from the Environmental Molecular Sciences Laboratory, a DOE Office of Science User Facility sponsored by the Biological and Environmental Research program under Contract No. DE-AC05-76RL01830. All work was conducted under the auspices of the US Department of Energy under Contract DE-AC52-07NA27344. LLNL-JRNL-2022143

