## Supplementary material for "Microbial communities in tropical soils are highly resilient to fluctuating redox conditions": Figure S1

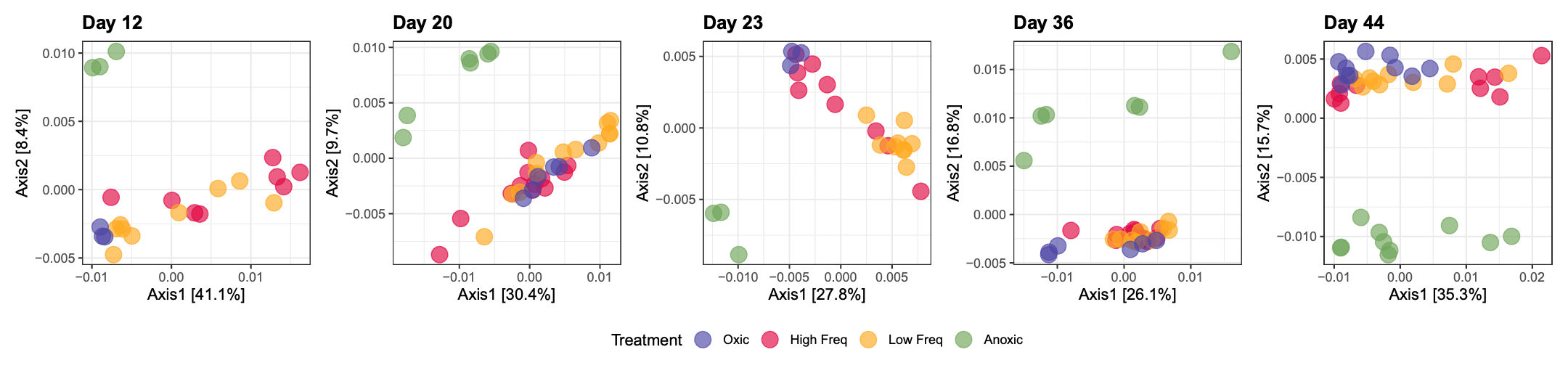

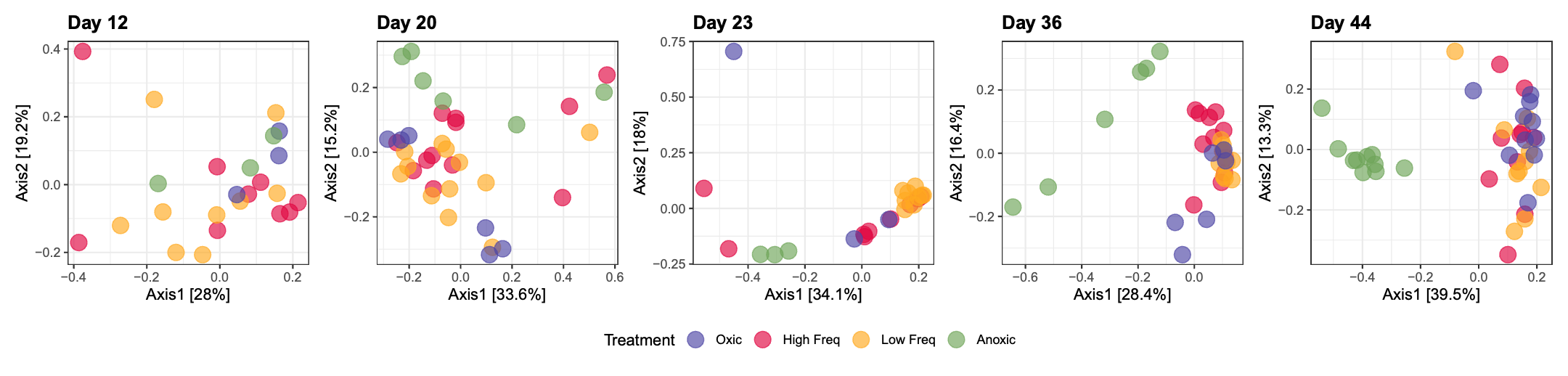

**Figure S1** Impacts of redox conditions on the soil bacterial community (top) and fungal community (bottom) over time.
