## Supplementary material for "Microbial communities in tropical soils are highly resilient to fluctuating redox conditions": Figure S2

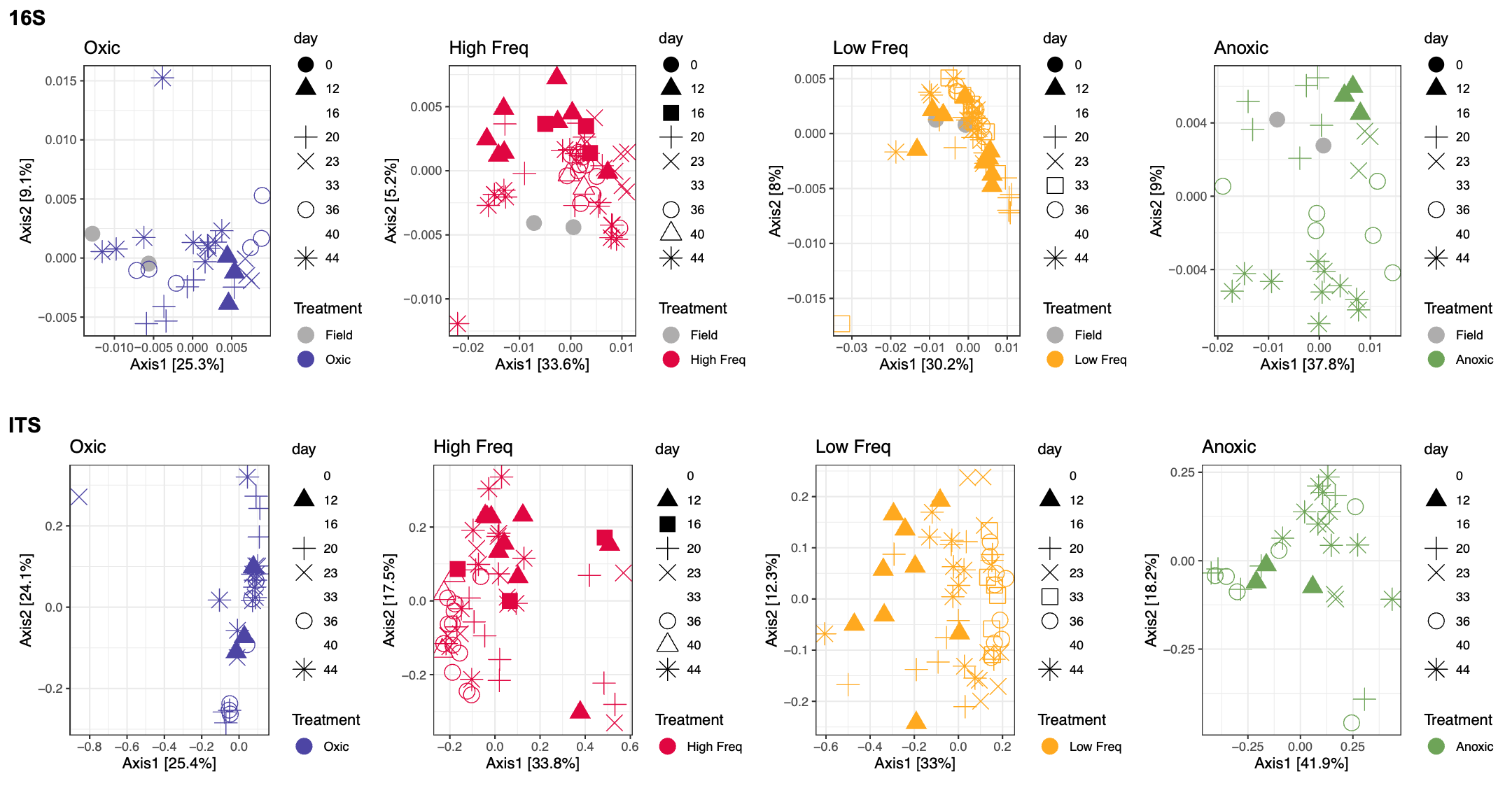

**Figure S2.**  Impact of each redox condition compared to the field composition for bacterial (top) and fungi (bottom).
