## Supplementary material for "Microbial communities in tropical soils are highly resilient to fluctuating redox conditions": Figure S3

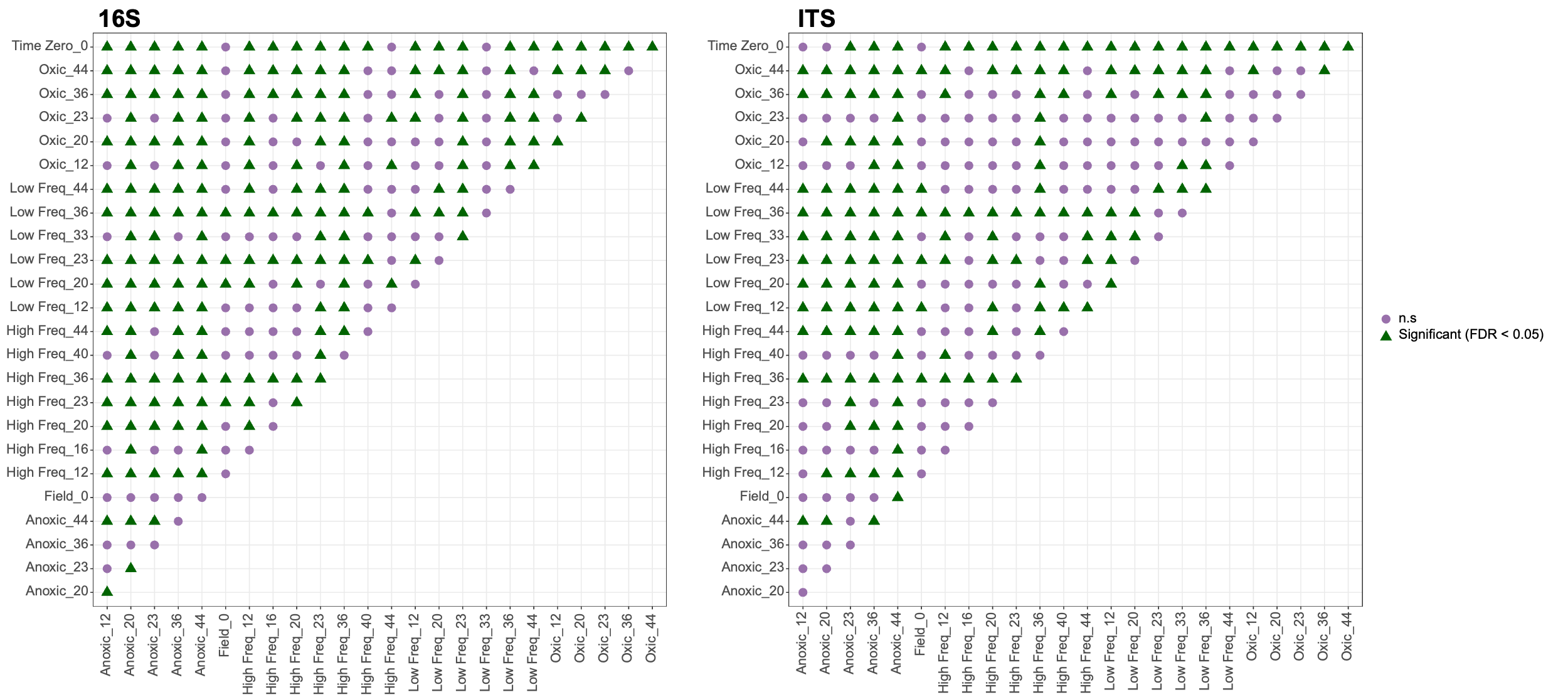

**Figure S3.**  Pairwise PERMANOVA with multiple comparison corrections for the bacterial community composition for each treatment_day. Significance denoted by green triangle (P ≤ 0.05).
