## Supplementary material for "Microbial communities in tropical soils are highly resilient to fluctuating redox conditions": Figure S4

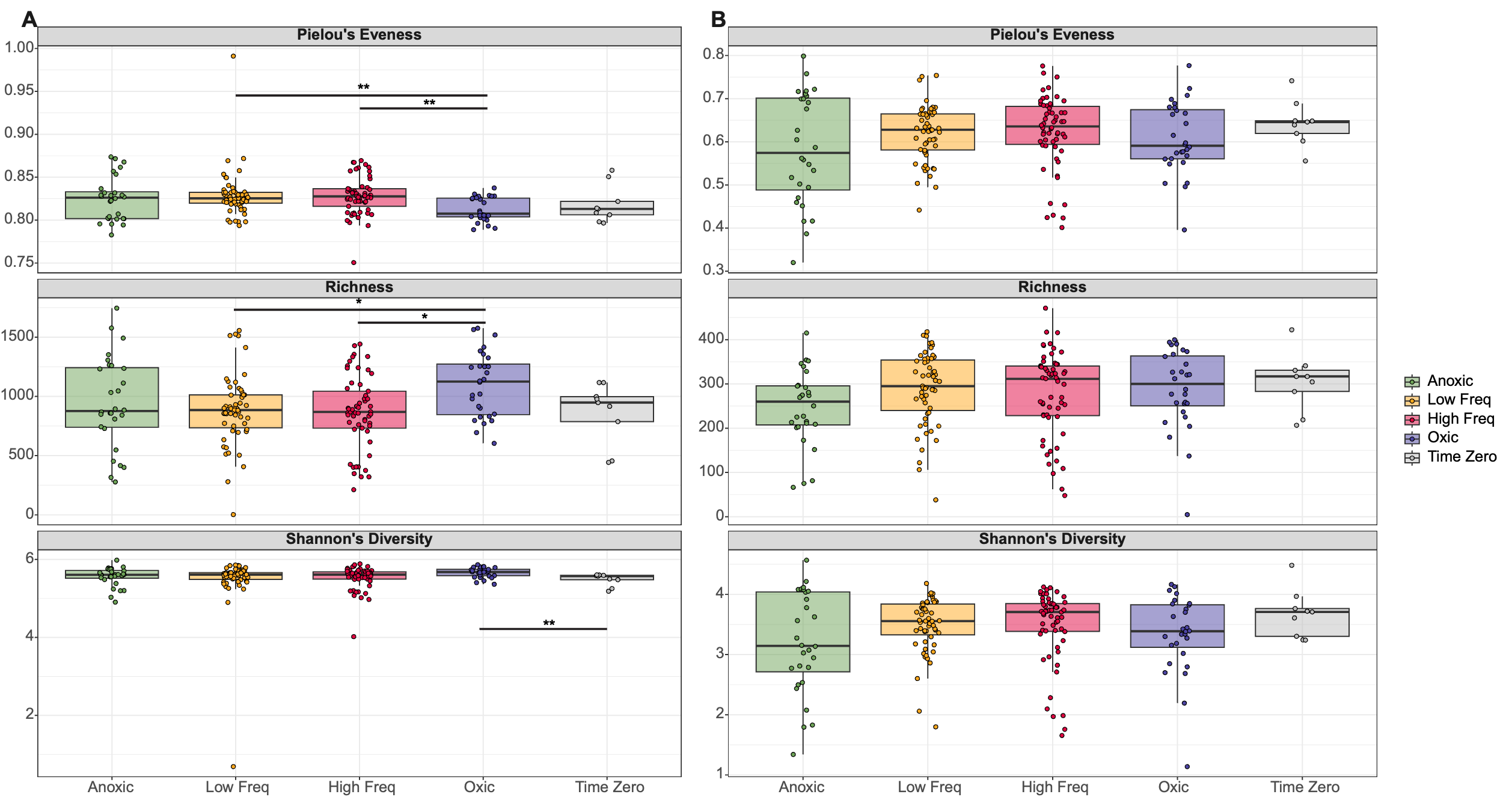

**Figure S4.** Bacterial community (A) and fungal community (B) alpha diversity. Control samples are T_0_ and Field condition. * p < 0.05, ** p ≤ 0.01
