## Supplementary material for "Microbial communities in tropical soils are highly resilient to fluctuating redox conditions": Figure S5

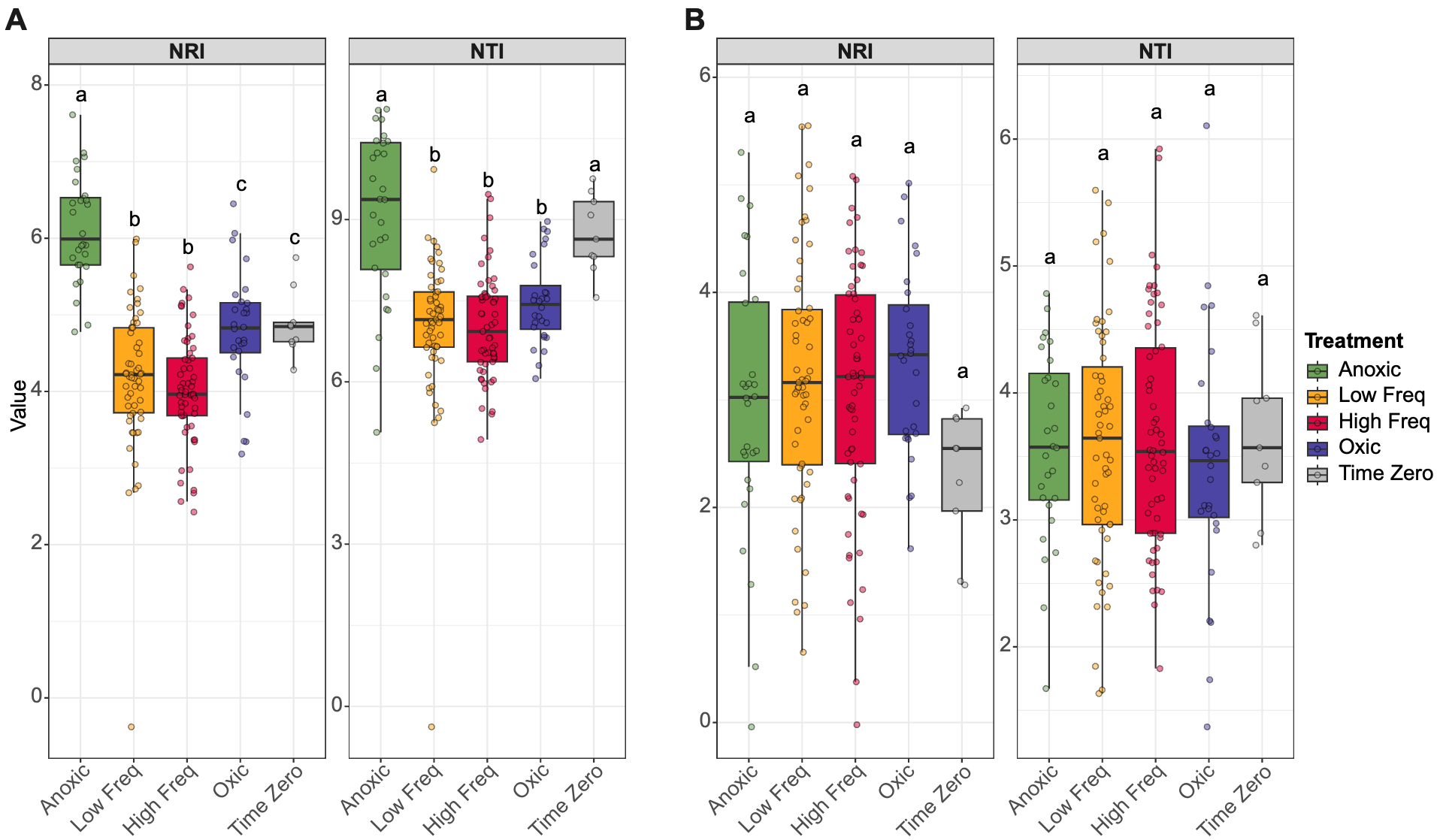

**Figure S5.** Bacterial community (A) and fungal community (B) NRI and NTI. Differing letters indicate significant differences (Dunn’s test or ANOVA-Tukey P < 0.05)
