## Supplementary material for "Microbial communities in tropical soils are highly resilient to fluctuating redox conditions": Figure S6

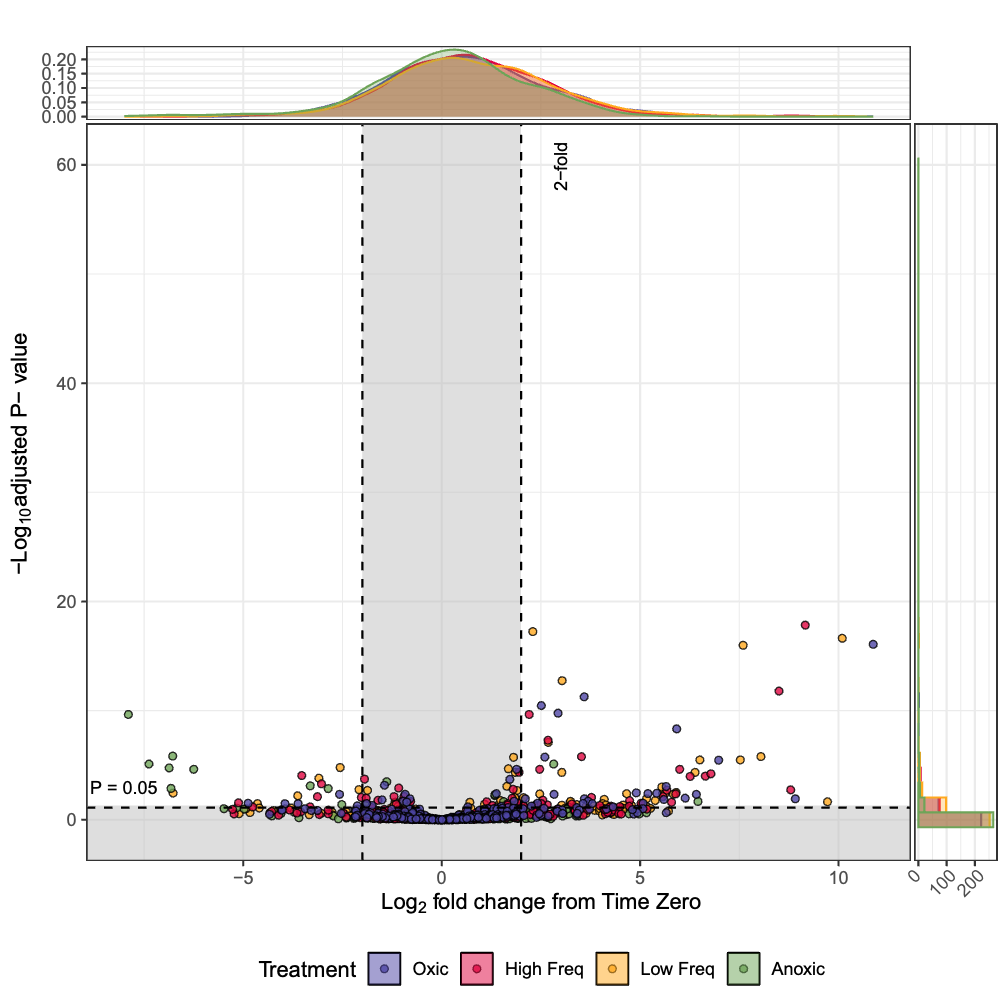

**Figure S6.**  Significant (unshaded plot area) and nonsignificant (shaded area) Log_2_ fold change response of fungal ASVs in each treatment (colors) relative to T0 identified using DESeq2. The vast majority of ASVs did not change relative to time zero (top) and/or did not pass false discovery rate of P = 0.05 (right). P-values are adjusted for multiple comparisons using ‘Benjamini-Hochberg’ correction.
