## Supplementary material for "Microbial communities in tropical soils are highly resilient to fluctuating redox conditions": Figure S7

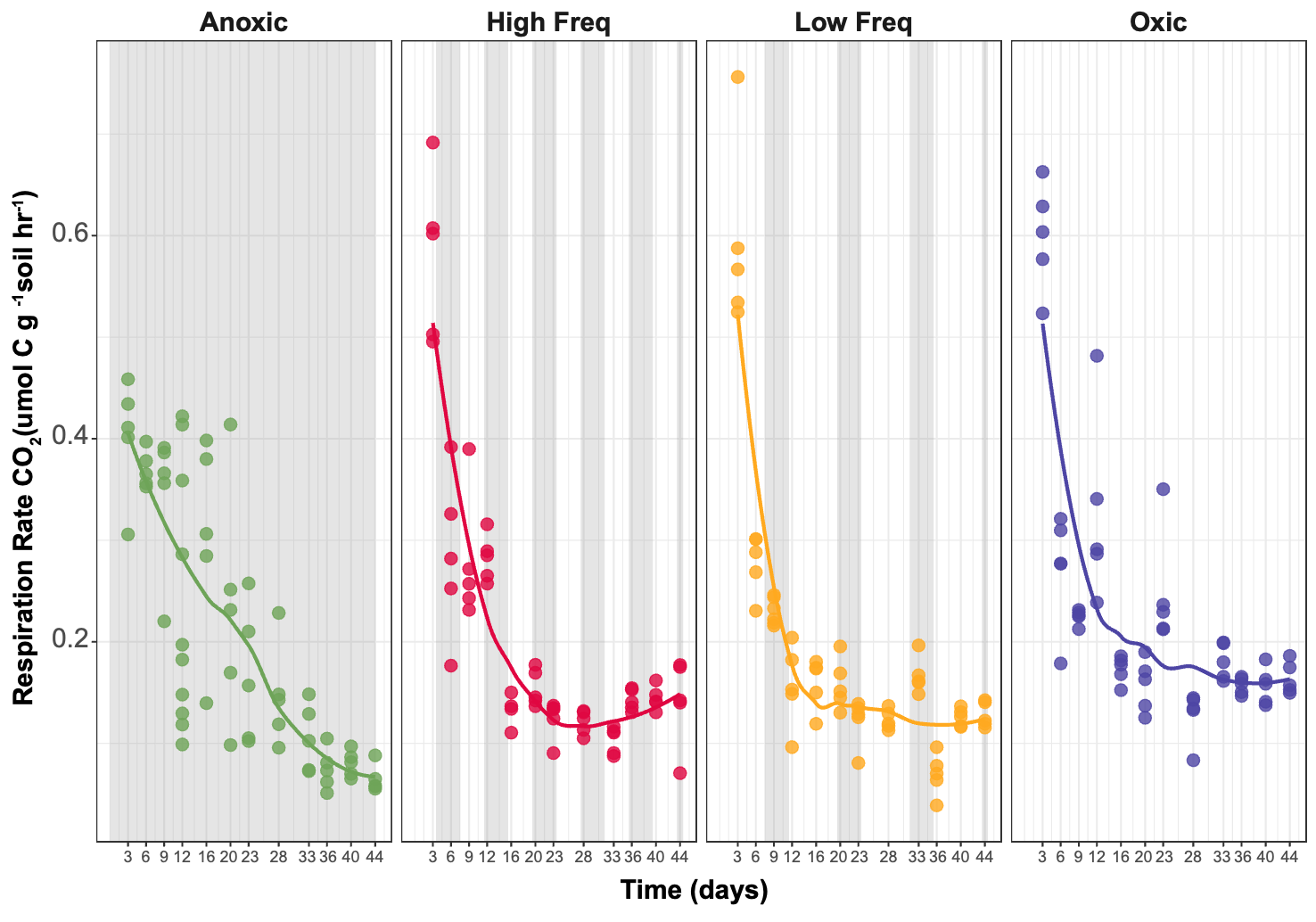

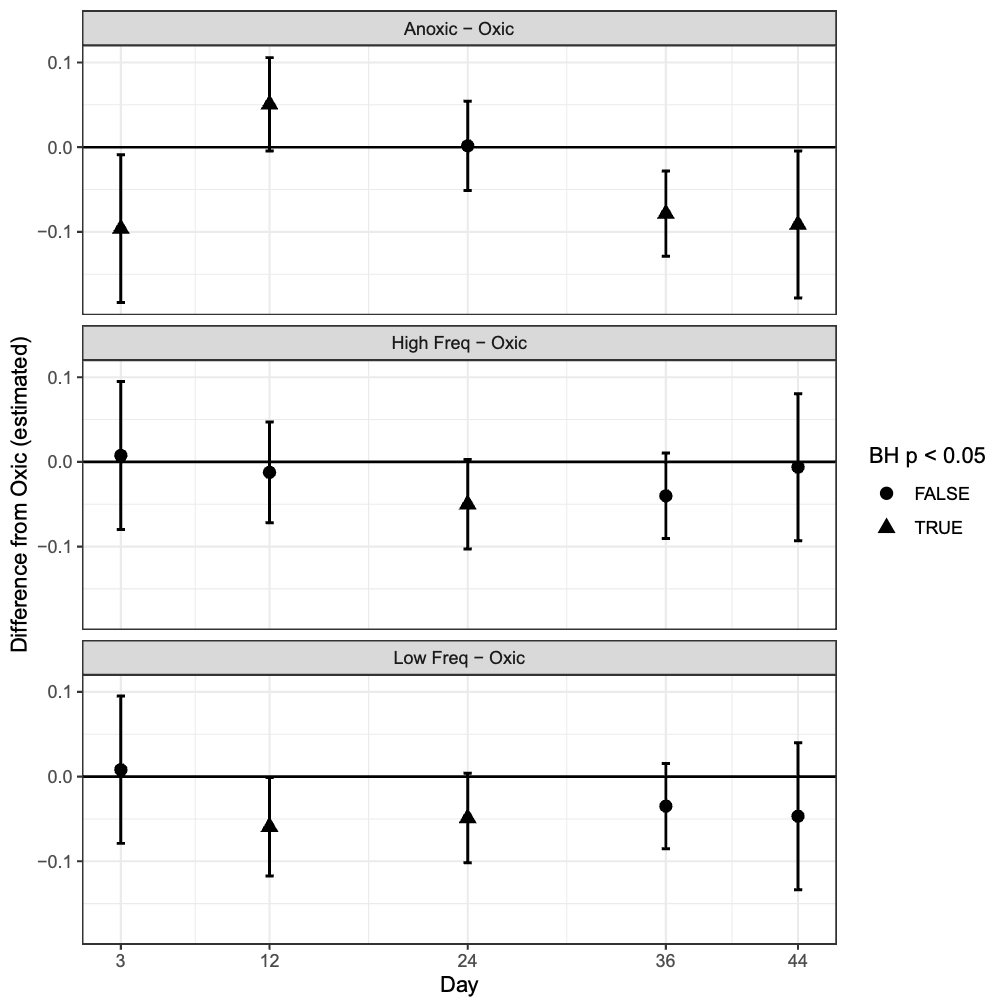

**Figure S7**. A) CO_2_ measurements over 44-day experimental period. Gray areas indicate anoxic periods and white areas oxic. B) Estimated marginal mean differences from mixed-effects modeling of CO_2_ flux over time. Significant differences from static oxic condition are denoted by triangles and error bars are 95% CIs. Points above 0 indicate higher CO_2_ flux compared to static oxic treatment and points below 0 indicate lower CO_2_ flux compared to static oxic.
