## Supplementary figures and images for "Microbial communities in tropical soils are highly resilient to fluctuating redox conditions"

### Table S1

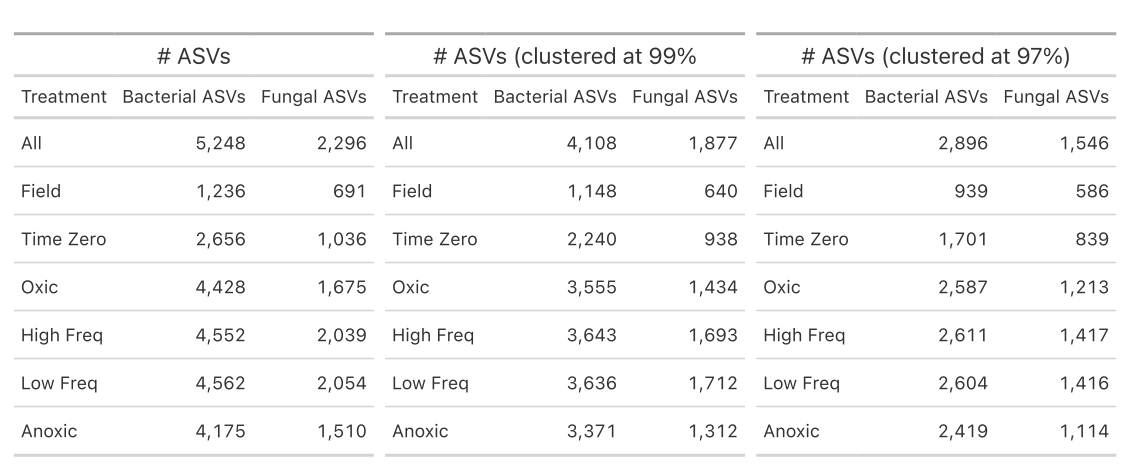

**Table S1.**  ASV counts per treatment.
