## Supplementary material for "Microbial communities in tropical soils are highly resilient to fluctuating redox conditions": Table S2

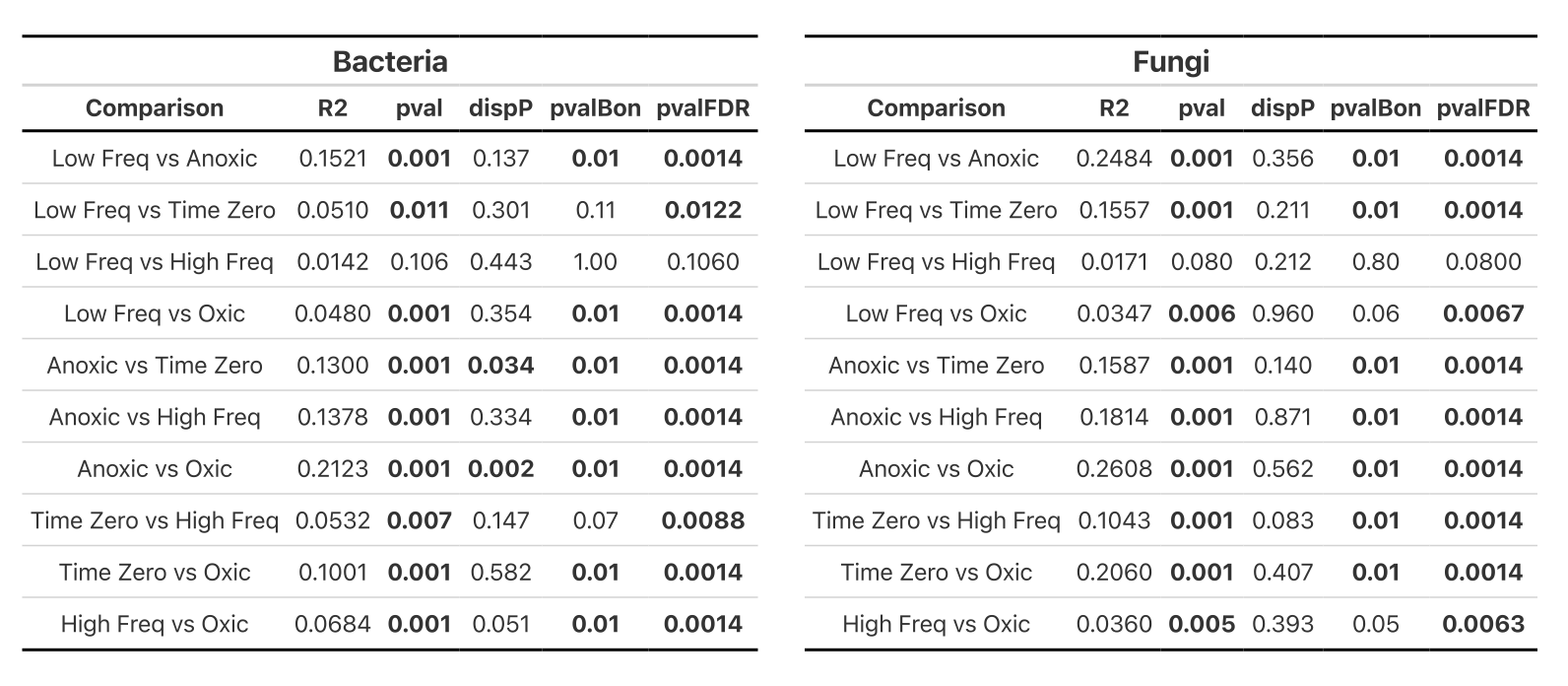

**Table S2.** Pairwise PERMANOVA with multiple comparison corrections for the bacterial (left) and fungal (right) community composition clustered at 97% for each treatment (includes full time series).
