## Supplementary material for "Microbial communities in tropical soils are highly resilient to fluctuating redox conditions": Table S3

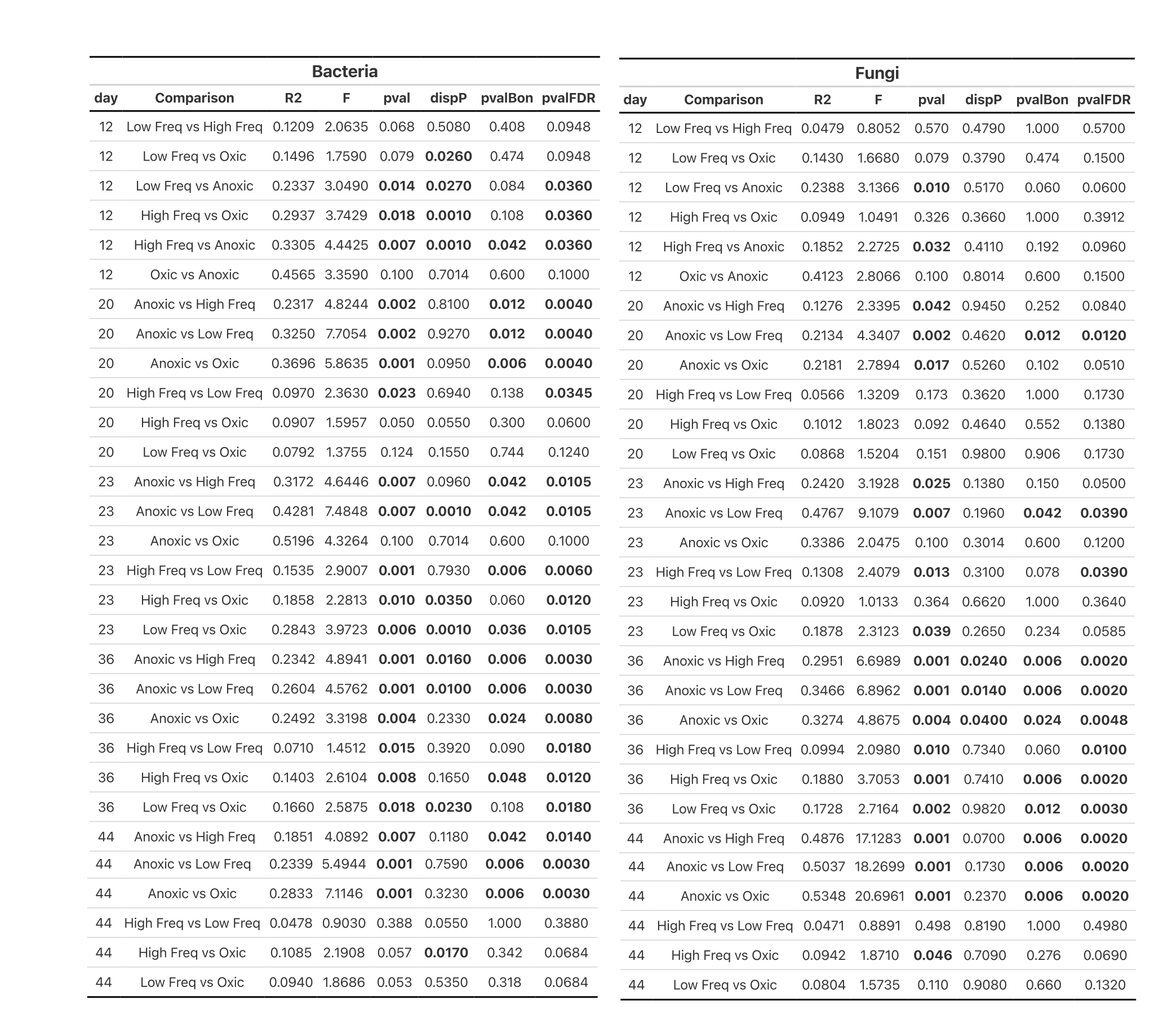

**Table S3.** Pairwise PERMANOVA with multiple comparison corrections for the bacterial community composition for each day with sampling from all 4 redox treatment (days 12, 20, 22, 36, 44).
