## Supplementary material for "Microbial communities in tropical soils are highly resilient to fluctuating redox conditions": Table S4

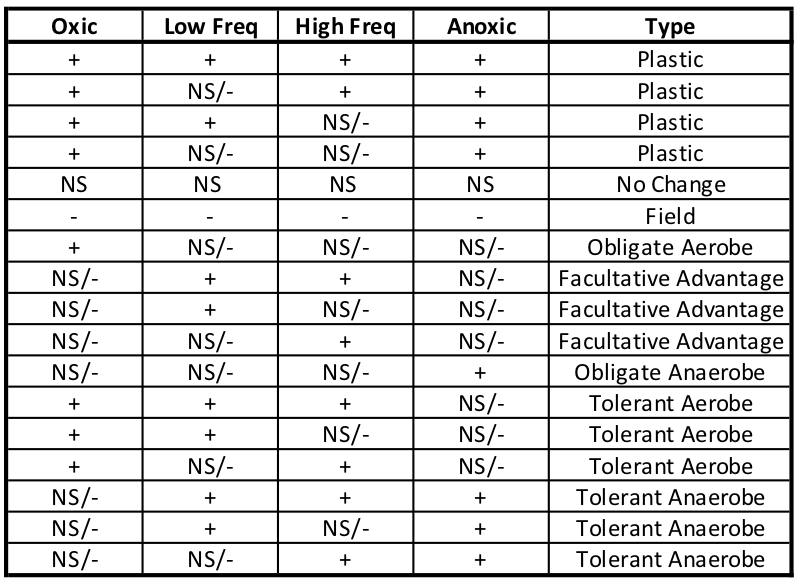

**Table S4.**  For each time point, each bacterial ASV was classified into a response type based on the pattern of how each ASV was impacted by the redox conditions collectively. Response of each ASV under each redox condition was determined using log2 fold change (with BH multiple comparisons correction) for that time point compared to T_0_. (+) significantly enriched, (-) significantly depleted, (NS) not significantly impacted.
