## Supplementary material for "Microbial communities in tropical soils are highly resilient to fluctuating redox conditions": Table S5

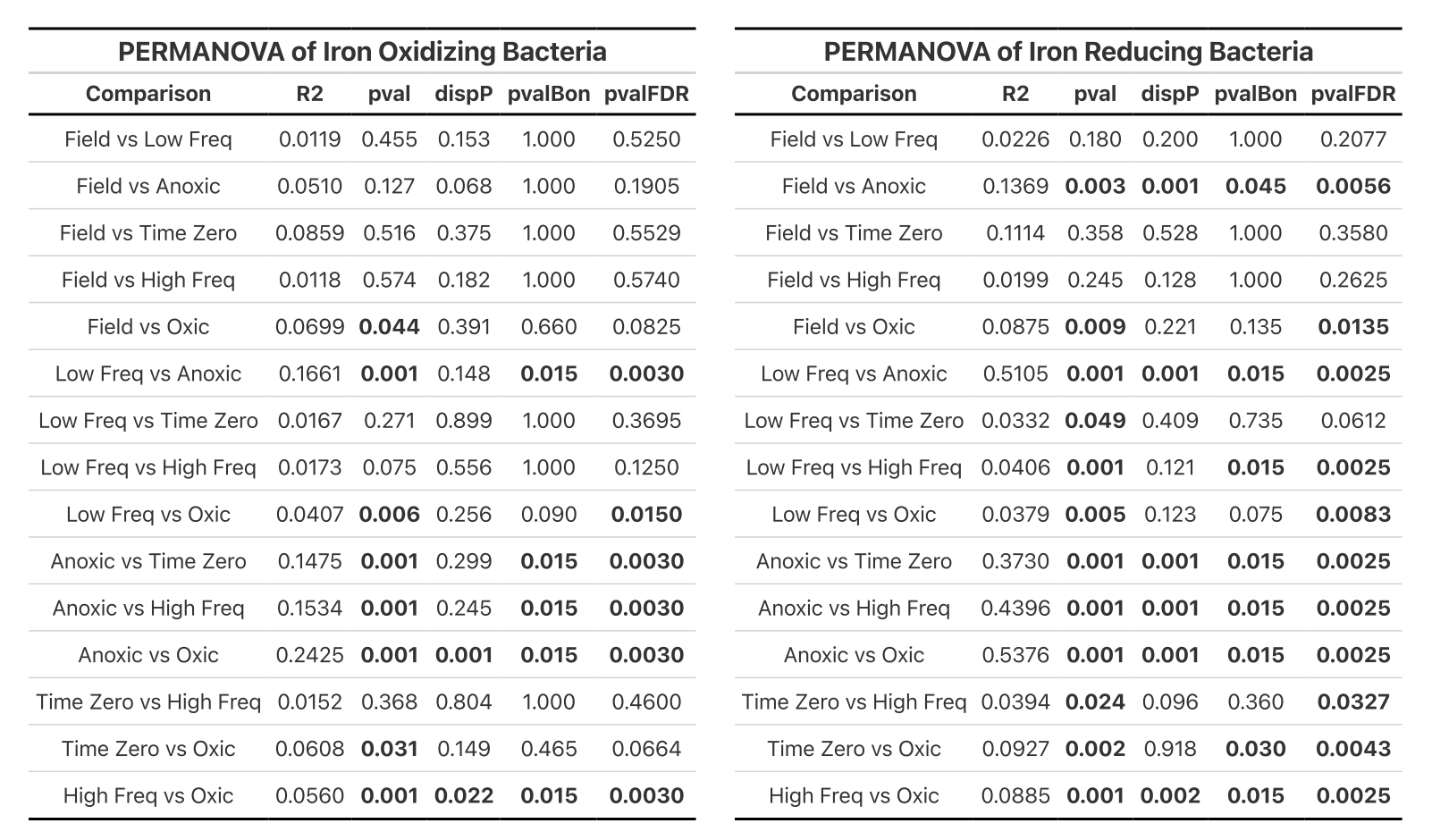

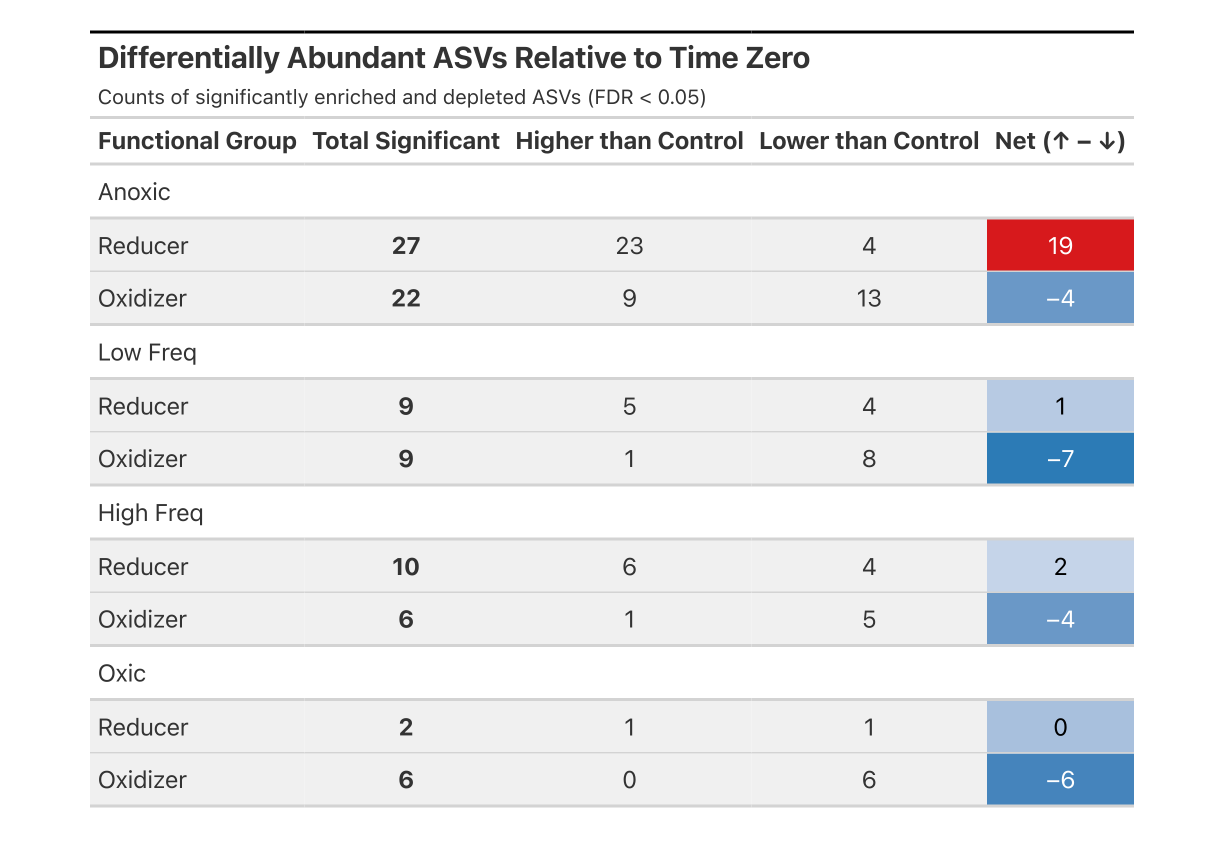

**Table S5. Iron cycling bacteria PERMANOVA and differential abundance**

A) Pairwise PERMANOVA results on iron cycling bacteria. B) Counts of differentially abundant ASVs relative to Control (Time Zero + Field) in each redox condition.
